# Gut archaea drive human MAIT cell responses

**DOI:** 10.64898/2026.09.11.750638

**Authors:** Natasha Fisher-Pearson, Martin J. Lett, Isaac W. D. Maylam, Connor J. S. Philp, Mariolina Salio, Richard J. Suckling, Robert Simmons, Keir Barnbrook, Benjamin Swift, Walter Jaoko, Freddie Kibengo, William Kilembe, Eduard J. Sanders, Mark C. Coles, Claire F. Pearson, Paola Cicconi, Tomáš Hanke, Nicholas M. Provine, Paul Klenerman, Jethro S. Johnson

**Author notes:** These authors contributed equally to this work (Natasha Fisher-Pearson, Martin J. Lett). These authors jointly supervised this work (Nicholas M. Provine, Paul Klenerman, Jethro S. Johnson).

## Abstract

Archaea, the third domain of life, are a gut microbiome constituent, but their role in shaping immunity is poorly understood. Mucosal-associated invariant T (MAIT) cells sense microbial metabolites through their T cell receptor and contribute to antimicrobial defence and barrier homeostasis. Key sources of the major MAIT cell antigen 5-OP-RU within the human gut microbiome are unknown. Analysing microbiomes from UK, Zambian, Kenyan and Ugandan donors, we found strong correlations between archaeal riboflavin biosynthesis and circulating MAIT cell frequency and activation. Stool 5-OP-RU levels correlated with archaeal abundance. Cultures of methanogenic archaea, including the human commensal *Methanobrevibacter smithii*, produced 5-OP-RU and strongly activated MAIT cells without inflammation. We demonstrate a critical role for archaea in shaping human immunity, with implications for microbiome-mediated regulation of health.

## Introduction

The human gut microbiome is well recognised for its role in shaping the immune system in both health and disease. However, as a significant proportion of the gut microbiome’s taxonomic and functional diversity remains poorly characterised, the specific features that drive many axes of microbe-immune interaction are not yet fully understood. This is particularly true for the relatively understudied non-bacterial domains of life such as archaea, whose relationship with human health and disease remains unclear (*1*).

Mucosal-associated invariant T (MAIT) cells are a key axis of microbe-immune communication (*2*). An abundant innate-like T cell subset with direct dependence on the microbiome, human MAIT cells are defined by their expression of a semi-invariant Vα7.2−Jα33/12/20 T cell receptor (TCR) (*3, 4*). The MAIT TCR primarily recognises water-soluble metabolic derivatives of microbial riboflavin biosynthesis (*5*), presented by the MHC class Ib molecule MR1. The most potent activating ligand is 5-(2-oxopropylideneamino)-5-d-ribitylaminouracil (5-OP-RU), generated by non-enzymatic condensation of the riboflavin biosynthesis intermediate 5-amino-6-d-ribitylaminouracil (5-A-RU) and methylglyoxal (*6*). 5-OP-RU can diffuse rapidly from the intestine to the blood (*7*) and is necessary and sufficient for MAIT cell development in rodents (*8, 9*). As the riboflavin biosynthesis pathway (RBP) is absent in animal cells, the specificity of the MAIT TCR enables distinction between host and microbe (*5, 10*). Extensive work in mouse models has linked the presence of the RBP in bacteria and fungi with MAIT cell function and development (*11*). In addition to activation via their TCR, MAIT cells can be activated by combinatorial cytokine signalling (*12, 13*). The balance of these two inputs tunes MAIT cell functionality (*14*), enabling MAIT cells to play diverse roles in antibacterial (*15*–*21*) and antiviral defence (*13*), tissue repair (*9, 22*) and maintenance of barrier homeostasis (*7, 23*).

Circulating MAIT cell abundance varies by up to 100-fold between healthy individuals (*24*), but the biological drivers of this variation – including the contribution of the gut microbiome – remain unclear. This knowledge gap reflects the difficulty of detecting and quantifying 5-A-RU/5-OP-RU at sub-nanomolar concentrations (*8, 25, 26*). Furthermore, almost all studies of human MAIT cells come from urban, developed societies, and it remains unknown how MAIT cell frequency, phenotype and functionality vary globally. Similarly, there is a bias in microbiome studies towards populations from developed countries (*27*). As human microbiomes show substantial inter-individual and inter-population variation (*28, 29*), we hypothesised that features of a complex human gut microbiome are a major driver of global variation in circulating MAIT cell abundance and activation. Notably, this includes features that may be lost in urban, Western populations, and that are not captured by next-generation sequencing approaches biased toward detection of bacteria.

Here we address this hypothesis by comparing MAIT cell abundance and activation across four African populations and one UK population. We show that circulating MAIT cells are less abundant but more activated in African populations compared to Europeans. Using shotgun metagenomic sequencing, we identify archaeal riboflavin synthesis as a major correlate of MAIT cell phenotype in humans. By developing a novel, sensitive, soluble TCR-based assay for MAIT antigen quantification, we show for the first time that archaea can produce MAIT cell ligands and activate MAIT cells *in vitro*, revealing a previously unrecognised pathway through which the human immune system senses the archaeome.

## Results

### Variation in MAIT cell abundance and phenotype across individuals and global populations

To characterise the differences in MAIT cell abundance and phenotype across individuals and diverse human populations, we performed spectral flow cytometry on peripheral blood mononuclear cells (PBMCs) from healthy volunteers from Oxford, UK (n = 10); Lusaka, Zambia (n = 22); Nairobi, Kenya (n = 22); Kilifi, Kenya (n = 22) and Masaka, Uganda (n = 22) (Fig. 1A; demographic information in table S1; gating strategies in Fig. S1 and Fig. S2). Individuals from the Oxford population had a significantly higher frequency and count of circulating MAIT cells than all other populations studied when correcting for age and sex (Fig. 1B and Fig. S3A). Males had significantly more abundant MAIT cells than females across the whole study population (Fig. S3, B and C). Other unconventional T cell subsets showed minimal geographic variation (Fig. S3, D to G).

**Fig. 1.**
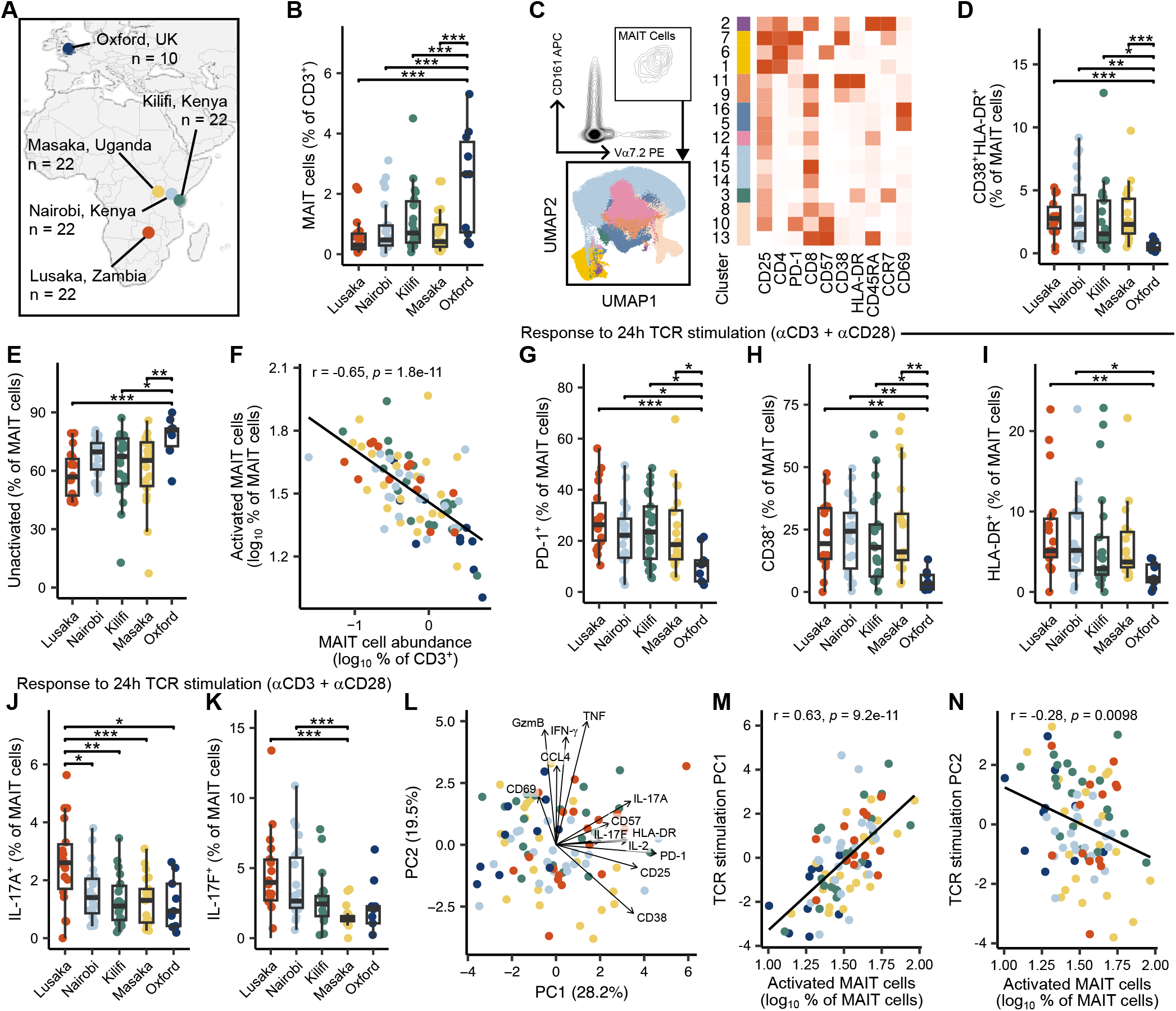
Circulating MAIT cells show inter-individual and inter-population variation in abundance, activation phenotype and functionality. (**A**) Sample number and location of study sites. (**B**) Abundance of MAIT cells by study site. Boxplots and individual data points show raw data distributions. P-values from linear model adjusting for sex and age. (**C**) UMAP and heatmap of MAIT cell activation clusters. (**D** and **E**) Abundance of MAIT cell clusters (**D**) 9 and 11 (CD38^+^HLA-DR^+^) and (**E**) 4, 14 and 15 (‘unactivated’) by study site. (**F**) MAIT abundance (percent of CD3^+^ T cells) by abundance of activated MAIT cells (percentage of clusters 4, 14 and 15 subtracted from 100). (**G** to **K**) Abundance of (**G**) PD-1^+^, (**H**) CD38^+^, (**I**) HLA-DR^+^, (**J**) IL-17A^+^ and (**K**) IL-17F^+^ MAIT cells following 24-hour anti-CD3 and anti-CD28 stimulation by study site. (**L**) PCA on z-score normalised change in percent expression of markers by MAIT cells following TCR stimulation. Arrows represent loadings. (**M** and **N**) Abundance of activated MAIT cells (percent of MAIT cells) by (**M**) PC1 and (**N**) PC2. All panels represent data from n = 92 biologically independent samples (Lusaka n = 18; Nairobi n = 22; Kilifi n = 21; Masaka n = 21; Oxford n = 10). Boxplots denote median, 25th and 75th percentiles, and whiskers to the largest value within 1.5 × interquartile range of the percentiles. Data distribution was assessed for normality using the Shapiro-Wilk test; significance was determined by ANOVA with Tukey’s post-hoc correction (two-sided) where assumptions met; otherwise, Kruskal-Wallis H test with Dunn’s post-hoc test (two-sided) with Bonferroni’s correction used. *, P < 0.05; **, P < 0.01; ***, P < 0.001 (adjusted p-values). For scatter plots, r denotes Pearson’s correlation coefficient. Data points are coloured by study site. Line represents the linear regression fit (least-squares line) across all data points.

We then performed unsupervised meta-clustering of gated MAIT cells according to expression of activation markers directly *ex vivo* (Fig. 1C). MAIT cells from populations in Lusaka, Nairobi, Kilifi and Masaka all exhibited a significantly higher proportion of CD38^+^HLA-DR^+^ activated MAIT cells compared to the Oxford population (Fig. 1D). Conversely, the proportion of MAIT cells not expressing any activation markers was significantly higher in the Oxford population compared to three African populations (Fig. 1E). The abundance of circulating MAIT cells showed a strong negative correlation with degree of activation (Fig. 1F) as previously observed (*2*).

To understand MAIT cell function, we stimulated PBMCs for 24 hours with either cytokine (interleukin [IL]-12 and IL-18) or TCR stimulation (anti-CD3 and anti-CD28 monoclonal antibodies) to capture the two modes of MAIT cell triggering. Following cytokine stimulation, there were no major inter-population differences in activation phenotype and cytokine production (Fig. S4). However, following TCR stimulation, a significantly lower percentage of MAIT cells expressed activation markers PD-1, CD38 and HLA-DR in the Oxford population compared to all other populations (Fig. 1, G to I). Furthermore, following TCR stimulation, there were significant inter-population differences in production of IL-17A and IL-17F (Fig. 1, J and K). Notably, there were no inter-population differences in the type 1 response (such as IFN-γ and granzyme B production) (Fig. S4). We performed a principal component analysis to summarise the response to TCR stimulation (Fig. 1L). This showed significant variation across populations ((R^2^ = 0.089, F = 2.13, p = 0.004, PERMANOVA). The first principal component – driven by expression of IL-17A, IL-17F, PD-1, CD38 and other activation markers – was significantly correlated with the percentage of activated MAIT cells stained directly *ex vivo* (Fig. 1M). By contrast, the second principal component – driven by expression of type 1 cytokines such as granzyme B, TNF and IFN-γ – was negatively correlated with MAIT activation directly *ex vivo* (Fig. 1N).

Overall, this suggested that African populations had a more activated MAIT phenotype when measured directly *ex vivo* and distinct response to TCR triggering compared to those in the UK. Furthermore, all populations exhibited substantial inter-individual variability in MAIT cell abundance, activation and functionality.

### Identifying microbiome correlates of MAIT cell phenotype

Next, we next explored whether differences in gut microbiome composition and functional potential could explain inter-individual variation in MAIT cell phenotype. We performed shotgun metagenomic sequencing on stool samples collected from the cohort. The faecal microbiomes of the Oxford cohort exhibited significantly lower species richness than several African populations (Fig. 2A). Besides their lower diversity, the microbiomes of the Oxford cohort showed a distinct genus-level composition to the African populations (R^2^ = 0.14, F = 3.86, p < 0.0001, PERMANOVA) (Fig. 2B and Fig. S5A). Microbiomes from the Oxford cohort were also distinct in their functional potential (abundance of MetaCyc metabolic pathways (*30*)) (R^2^ = 0.22, F = 6.93, p < 0.0001, PERMANOVA) (Fig. 2C).

**Fig. 2.**
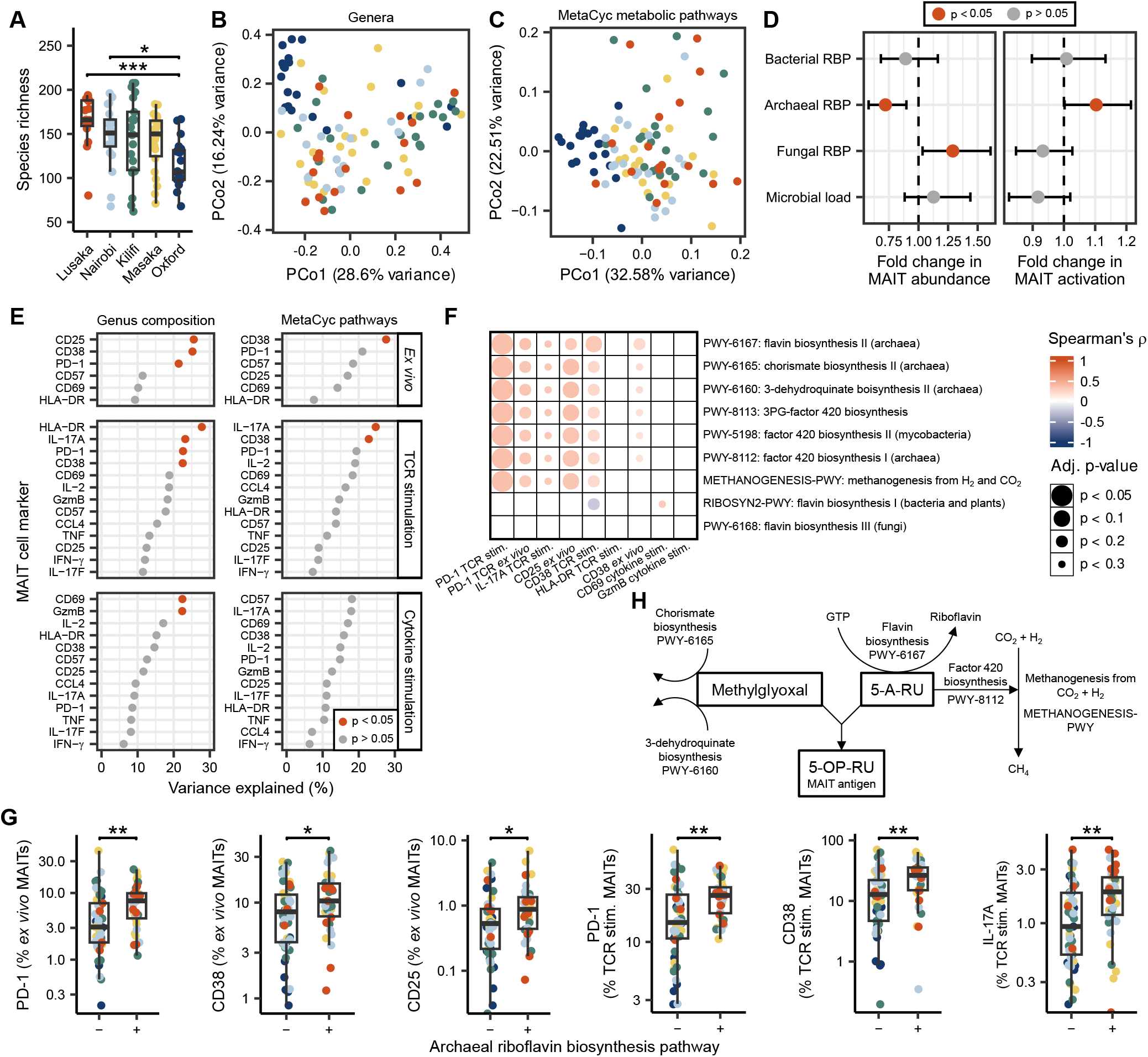
Archaeal riboflavin biosynthesis in the microbiome is correlated with MAIT cell phenotype and functionality. (**A**) Species richness by study site. Normality assessed by Shapiro-Wilk test. Significance assessed by ANOVA with Tukey’s post-hoc correction (two-sided). *, p < 0.05; **, p < 0.01; ***, p < 0.001 (adjusted p-values). (**B**) PCoA on Bray-Curtis dissimilarity (BCD) of genus-level relative abundances. Study site is colour coded. (**C**) PCoA on BCD of abundance of MetaCyc gene pathways. (**D**) Coefficient plot of microbiome predictors of log-transformed MAIT abundance and log-transformed abundance of activated MAIT cells. Points indicate estimated regression coefficients (β) from a linear model including z-scaled abundance of the bacterial, fungal and archaeal riboflavin synthesis pathways, z-scaled microbial load, sex and age with study site as a random effect. Bars indicate 95% confidence intervals. Colour denotes significance (p < 0.05) by type III F-tests with Satterthwaite’s approximation. (**E**) Variance explained (R-squared) from linear models (MAIT cell phenotypic marker ∼ PC1 + … + PC10), where PC1-PC10 are principal coordinates from a PCoA on genus- or pathway-level BCD. Colour denotes significance (p < 0.05) by overall model F-test. (**F**) Spearman’s rank correlations between MetaCyc gene pathways and MAIT cell phenotypic markers. Colour indicates Spearman’s ρ; circle size denotes Benjamini-Hochberg-adjusted p-values. (**G**) Abundance of MAIT cells expressing phenotypic markers (*ex vivo* or following TCR stimulation) in individuals where the archaeal riboflavin biosynthesis pathway (RBP) is detectable (+) or undetectable (−) by shotgun metagenomic sequencing. Significance determined by two-sided Mann-Whitney U test. (**H**) Schematic of MAIT cell antigen synthesis during archaeal methanogenesis. Figures [(**A**) to (**C**)] represent data from n = 100 biologically independent faecal samples (Lusaka n = 17; Nairobi n = 20; Kilifi n = 21; Masaka n = 22; Oxford n = 20). Figures [(**D**) to (**G**)] represent data from n = 84 biologically independent matched faeces and PBMCs (Lusaka n = 13; Nairobi n = 20; Kilifi n = 20; Masaka n = 21; Oxford n = 10). Boxplots denote median, 25th and 75th percentiles, and whiskers to the largest value within 1.5 × interquartile range of the percentiles.

We then investigated whether MAIT cell abundance and *ex vivo* activation phenotype could be predicted from abundance of riboflavin biosynthesis pathways (RBPs) in the microbiome using mixed effects linear models. Abundance of the archaeal RBP was significantly associated with MAIT cell abundance and activation, whereas abundance of the bacterial RBP was not a significant correlate (Fig. 2D) – a highly unexpected finding as archaea have never been shown to activate MAIT cells before. Abundance and prevalence of archaea was significantly higher in the populations from Lusaka and Masaka compared to Oxford (Fig. S5B). Using a likelihood ratio test, we found that the inclusion of microbial predictors significantly improved performance compared to a reduced model predicting MAIT abundance from age, sex and study site alone (p = 0.02).

Next, we investigated which MAIT cell phenotypic markers were significantly associated with microbiome structure. We leveraged individual linear models, with the first 10 principal coordinates of Bray-Curtis dissimilarity on genus-level composition (Fig. S5C) or pathway-level composition (Fig. S5D) as predictors, and percentage expression of a given MAIT cell phenotypic marker as the outcome. Microbiome genus- and pathway-level composition explained up to 27.8% of the variation in MAIT cell marker expression (Fig. 2E), with expression of HLA-DR, CD25, CD38, IL-17A, PD-1, CD69 and granzyme B significantly associated with microbiome structure (Fig. 2E).

To determine specifically which microbiome features were associated with variation in these markers of MAIT cell activation and function, we performed Spearman’s correlations between all MetaCyc pathways detected in the microbiome and percentage expression of markers identified in Fig. 2E. Out of all 476 MetaCyc pathways, archaeal riboflavin biosynthesis was the strongest positive correlate of PD-1 expression *ex vivo* and second strongest correlate of IL-17A production following TCR stimulation (Fig. 2F). The abundance of MAIT cells expressing PD-1, CD38 and CD25 *ex vivo* and PD-1, CD38 and IL-17A following TCR stimulation was significantly higher in individuals where the archaeal RBP was detectable (Fig. 2G). A cluster of related archaeal pathways were also in the top 15 most strongly positively correlated pathways with CD25, PD-1 and IL-17A expression, including chorismate biosynthesis, 3-dehydroquinate biosynthesis, factor 420 biosynthesis and hydrogenotrophic methanogenesis (Fig. 2F). Chorismate biosynthesis and 3-hydroquinate biosynthesis generate methylglyoxal, which spontaneously reacts with 5-A-RU generated during riboflavin synthesis to form the MAIT cell antigen 5-OP-RU (summarised in Fig. 2H). 5-A-RU is the precursor for factor 420, an essential cofactor for hydrogenotrophic methanogenesis in archaea. By contrast, flavin biosynthesis pathways in bacteria and fungi had minimal association with MAIT cell function and phenotype (Fig. 2F).

These data suggest that archaeal riboflavin biosynthesis – as an intermediate process within methanogenesis – in the microbiome is a significant predictor of MAIT cell phenotype and function in humans.

### A novel assay to quantify MAIT cell ligands

To determine whether archaeal riboflavin synthesis was a significant driver of 5-OP-RU production by the microbiome, we required a sensitive method for quantifying 5-OP-RU in human faeces. Existing cell-based bioassays are susceptible to modulation by pro-inflammatory stimuli, whereas conventional physicochemical approaches lack sensitivity (*25*). We therefore developed a novel cell-based assay in which THP-1 cells engineered to over-express MR1 are incubated with either defined concentrations of synthetic 5-OP-RU or biological samples of interest. Following antigen loading, cells are stained with a soluble MAIT TCR optimised to exhibit high affinity and selectivity for MR1/5-OP-RU complexes. The combination of MR1-mediated antigen capture and TCR-based detection provided two independent layers of specificity, enabling sensitive quantification of bioactive 5-OP-RU in the picomolar range (Fig. 3A and Fig. S6). To evaluate the specificity of the assay, we analysed supernatants from bacterial monocultures. No detectable signal was observed for supernatants from cultured *Enterococcus faecalis* and the Δ*ribA* mutant of the laboratory *Escherichia coli* strain BL21 (which cannot synthesise 5-OP-RU). By contrast, a clear signal was detected in wild-type BL21 supernatants, irrespective of whether the growth medium was supplemented with riboflavin. This supports the specificity of the assay for microbially derived, riboflavin-pathway-dependent MAIT-cell antigens (Fig. 3B).

**Fig. 3.**
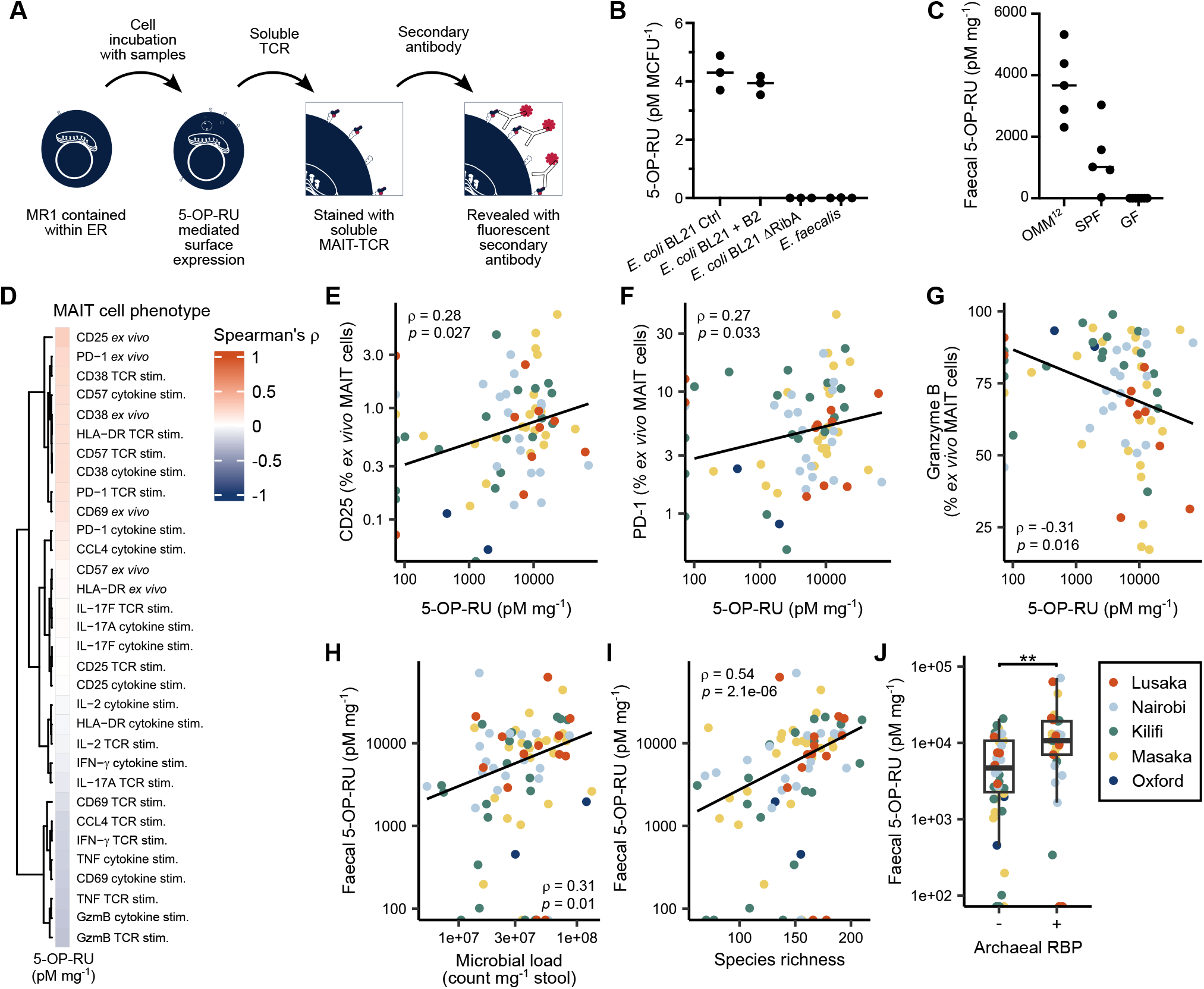
Faecal 5-OP-RU concentration correlates with circulating MAIT cell phenotype and archaeal riboflavin biosynthesis. (**A**) Schematic of assay used to quantify 5-OP-RU in biological samples. THP-1 cells expressing high levels of human MR1 were incubated for 4 h with either known concentrations of 5-OP-RU (standards) or test samples. Surface MR1– 5-OP-RU complexes were detected using a high-affinity engineered MAIT TCR fused to rabbit Fc, followed by staining with secondary antibodies and flow cytometric analysis. A standard curve relating pooled mean fluorescence intensity (MFI) to 5-OP-RU concentration was used to quantify 5-OP-RU in unknown samples. (**B**) Validation of assay specificity using culture supernatants from *Enterococcus faecalis*, wild-type BL21 *Escherichia coli* (with or without riboflavin [“B2”] supplemented) and *ribA*-deficient (*ΔribA*) BL21 *E. coli*. MCFU, million colony forming units. (**C**) Validation of assay specificity using faeces from germ-free, specific pathogen-free (SPF), or oligo-mouse microbiota 12 (OMM^12^)-colonised mice. (**D**) Heatmap of Spearman’s rank correlations between faecal 5-OP-RU concentrations and MAIT cell phenotypic or functional parameters measured directly *ex vivo* or following *ex vivo* stimulation. (**E** to **G**) Correlation between faecal 5-OP-RU concentration and circulating MAIT cell expression of (**E**) CD25 and (**F**) PD-1 directly *ex vivo* and (**G**) granzyme B following *ex vivo* TCR stimulation. (**H** and **I**) Correlation between faecal 5-OP-RU concentration and (**H**) faecal microbial load and (**I**) faecal microbiome species richness. (**J**) Faecal 5-OP-RU concentration in individuals with the archaeal riboflavin biosynthesis pathway (RBP) detectable (+) or undetectable (−) by shotgun metagenomic sequencing. Figures [(**E**) to (**I**)], ρ denotes Spearman’s correlation coefficient. Data points are coloured by study site. Line represents the linear regression fit (least-squares line) across all data points. (**J**) significance determined by two-sided Mann-Whitney U test. Boxplot denotes median, 25th and 75th percentiles, and whiskers to the largest value within 1.5 × interquartile range of the percentiles. Individual data points are shown. Figures [(**D**) to (**J**)] represent data from n = 74 (Lusaka n = 13; Nairobi n = 18; Kilifi n = 19; Masaka n = 22; Oxford n = 2) biologically independent faecal samples.

To further validate assay specificity, we quantified 5-OP-RU in faeces sampled from mice with two distinct microbiome complexities (specific pathogen-free [SPF] and oligo mouse microbiota 12 [OMM^12^]) mice alongside germ-free mouse controls. As expected, no detectable signal was observed in germ-free mouse faeces (Fig. 3C). SPF and OMM^12^ faeces exhibited distinct concentrations of 5-OP-RU, indicating that the assay was sensitive to differences in antigen-producing potential of the microbiota.

We then applied the assay to faeces from our human study population. We performed an exploratory correlative analysis to identify whether production of MAIT cell antigens by the microbiota was associated with expression of activation markers and production of cytokines by circulating MAIT cells (Fig. 3D). Faecal 5-OP-RU concentration positively correlated with expression of CD25 (Fig. 3E) and PD-1 (Fig. 3F) on circulating MAIT cells, suggesting repeated low-level activation *in vivo*. These two markers were identified as significantly positively correlated with the archaeal RBP in Fig. 2F. Faecal 5-OP-RU concentration was inversely associated with granzyme B production following TCR stimulation (Fig. 3G). This potentially reflects a distinct transcriptional program in chronically stimulated MAIT cells, as suggested by the negative association between *ex vivo* MAIT cell activation and type 1 functionality observed in Fig. 1N.

Next, we explored which microbiome features were associated with faecal 5-OP-RU concentrations. Faecal 5-OP-RU was positively correlated with both microbial load (Fig. 3H) and species richness (Fig. 3I). Faecal 5-OP-RU concentration was significantly higher for microbiomes where the archaeal riboflavin biosynthesis pathway was detectable by shotgun metagenomic sequencing (Fig. 3J). By contrast, there was no association between faecal 5-OP-RU concentration and abundance of the bacterial (ρ = -0.18, p = 0.16) or fungal (ρ = 0.18, p = 0.14) riboflavin biosynthesis pathways.

In case the associations between the archaeal riboflavin biosynthesis pathway, faecal 5-OP-RU concentration and MAIT cell phenotype were driven solely by differences between the European and African populations, we repeated key analyses with the UK population excluded. The associations remained significant (Fig. S7). Overall, this correlative evidence suggests that archaea represent a major source of MAIT cell antigens *in vivo* despite their low abundance in the microbiome and thus shape the phenotype and functionality of the MAIT cell compartment.

### Archaea produce MAIT cell ligands and activate MAIT cells *in vitro*

We next sought to experimentally validate whether archaea can produce MAIT cell ligands and directly activate MAIT cells. *Methanobrevibacter smithii* (the most prevalent and abundant archaeal species in the human gut) and a panel of bovine-associated *Methanobrevibacter* species, an environmental *Methanobacterium* species, and human pathogenic and commensal bacterial isolates were mono-cultured then 5-OP-RU was quantified in the culture supernatant using the assay described above. *M. smithii* produced high concentrations of 5-OP-RU relative to other species tested (Fig. 4A). Because methylglyoxal required for conversion of 5-A-RU to 5-OP-RU may be limiting in these culture conditions, MAIT cell antigens were quantified in culture supernatant supplemented with 0.1% methylglyoxal. *M. smithii* ranked among the strongest 5-A-RU producers, with a higher concentration in the supernatant than *Akkermansia muciniphila, Clostridium innocuum* and *Salmonella enterica* (Fig. 4A).

**Fig. 4.**
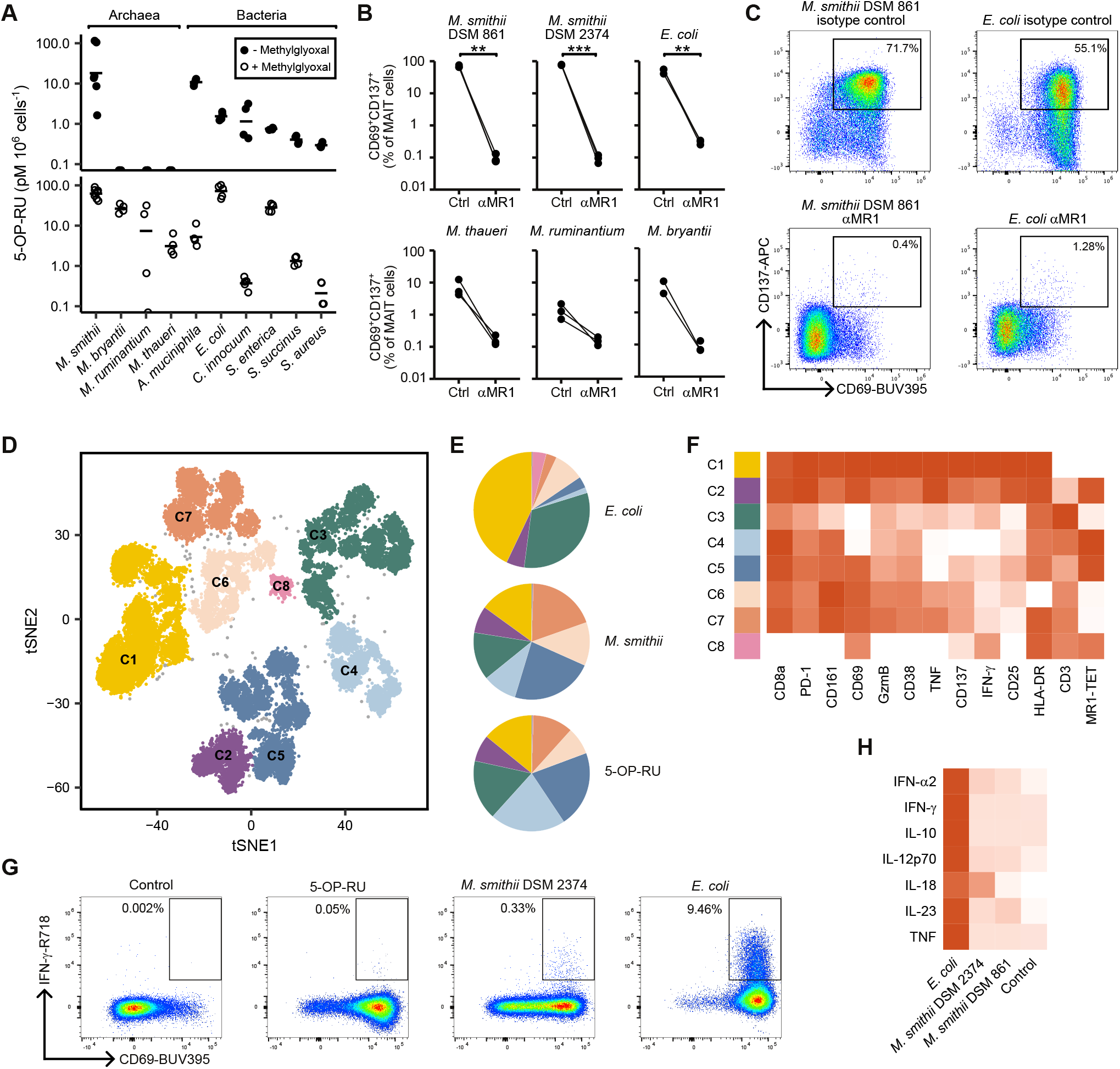
*Methanobrevibacter smithii* produces MAIT cell antigens and activates MAIT cells *in vitro*. (**A**) Concentration of 5-OP-RU in the culture supernatants of commensal and pathogenic microorganisms without supplementation or following supplementation with 0.1% methylglyoxal. *M. smithii, Methanobrevibacter smithii*; *M. bryantii, Methanobacterium bryantii*; *M. ruminantium, Methanobrevibacter ruminantium, A. muciniphila, Akkermansia muciniphila*; *E. coli, Escherichia coli*; *C. innocuum, Clostridium innocuum*; *S. enterica, Salmonella enterica*; *S. succinus, Staphylococcus succinus*; *S. aureus, Staphylococcus aureus*. (**B**) Expression of activation markers CD69 and CD137 on MAIT cells after 24 h stimulation with autologous monocytes exposed to the indicated microorganisms (MOI = 20), in the presence of an MR1-blocking antibody (αMR1) or an isotype control antibody (ctrl). Significance tested by two-sided t-test. (**C**) Representative flow cytometry plots depicting expression of CD69 and CD137 on MAIT cells after 24 h stimulation as in (**B**). (**D**) t-SNE projection of MAIT cells from four healthy donors after 36 h stimulation with autologous monocytes exposed to 5-OP-RU (2 nM), *Escherichia coli* or *Methanobrevibacter smithii* (MOI = 20), coloured by clusters identified by DBSCAN analysis. (**E**) Pie charts of distribution of cells from each stimulation condition within each cluster. (**F**) Heatmap of expression of functional and phenotypic markers (z-scaled MFI) across clusters. (**G**) Representative flow cytometry plots of CD69 and IFN-γ expression by MAIT cells from one donor included in the t-SNE analysis. (**H**) Heatmap showing normalised concentrations (pg/mL) of MAIT cell-associated cytokines in culture supernatants after 24 h stimulation with autologous monocytes and the indicated microorganisms in the presence of MR1-blocking antibody. Figure (**A**) represents data from n = 4 independent cultures per organism. Figures (**B**) represents PBMCs from n = 3 independent donors and [(**D**) to (**F**)] represent PBMCs from n = 4 independent donors.

We then assessed whether *M. smithii* could activate MAIT cells by stimulating PBMCs for 24 hours with fixed microbial cells, using autologous monocytes as antigen presenting cells (gating strategy in Fig. S8A). Two distinct strains of *M. smithii* induced strong upregulation of CD69 and CD137 on MAIT cells, with this response abolished by MR1 blockade (Fig. 4, B and C). A similar MR1-dependent activation pattern was observed with *Escherichia coli* and, to a lesser extent, the other *Methanobrevibacter* and *Methanobacterium* species.

Although CD69 and CD137 upregulation on MAIT cells were broadly comparable between *E. coli*- and *M. smithii*-stimulated PBMCs, cytokine secretion was markedly weaker in response to the archaea (Fig. S8B). Because MAIT cell activation is strongly influenced by TCR-independent cytokine signals, we hypothesised that the difference in activated MAIT cell functionality reflected limited engagement of innate immune receptors by archaea, resulting in reduced co-stimulation by monocytes. We therefore performed a 36-hour co-culture with *M. smithii, E. coli* or a 2 nM 5-OP-RU condition as a reference for isolated TCR stimulation. A t-SNE analysis with DBSCAN clustering identified eight distinct clusters based on activation marker and cytokine expression across stimulated MAIT cells (Fig. 4D). *M. smithii* and 5-OP-RU stimulation induced a similar distribution of MAIT cell clusters, which was distinct from the distribution induced by *E. coli* (Fig. 4E). *E. coli* stimulation induced a larger fraction of MAIT cells co-expressing CD69, CD137, TNF, IFN-γ and granzyme B, whereas both *M. smithii* and 5-OP-RU predominantly induced MAIT cells that produced neither IFN-γ nor TNF (Fig. 4, E and F). IL-17A and IL-17F-secreting MAIT cells were rare in this experiment and showed no significant differences between conditions, likely due to the short activation time (Fig. S8, C and D). Conventional gating approaches confirmed these observations (Fig. 4G).

To further investigate whether *M. smithii* induces inflammatory co-stimulation by monocytes, we analysed cytokine concentrations in supernatants from activated PBMCs from six independent donors. *E. coli* induced robust production of key MAIT-activating cytokines IL-12, IL-18, IL-23 and IFN-α in the presence of both MR1 blockade and isotype control (Fig. 4H and Fig. S8F). This response was not detected for either of the two *M. smithii* strains, further indicating that methanogenic archaea activate MAIT cells purely via the MAIT cell TCR, with minimal cytokine co-stimulation usually observed during bacterial activation.

## Discussion

In this study, we demonstrated that variation in the human gut microbiome is associated with the frequency and phenotype of circulating MAIT cells, an abundant innate-like T cell population with broad functionality in antimicrobial responses, tissue healing and immune homeostasis (*2*). By combining pathway-level microbiome analysis with functional antigen quantification and immune phenotyping, we identified microbiome configurations associated with increased local availability of 5-OP-RU and distinct circulating MAIT cell phenotypes. Unexpectedly, these analyses pointed to archaeal metabolism as a key source of MAIT antigens and not the bacterial species that comprise most of the gut microbiome. We further showed that methanogenic archaea can generate 5-A-RU and 5-OP-RU *in vitro* and strongly activate MAIT cells in a MR1-dependent manner.

This finding was enabled by the exploratory and geographically diverse design of our study. Although gut microbiome composition varies substantially across human populations (*28, 29*), most microbiome studies, and most studies on MAIT cells in health and disease, have focused on individuals from highly developed countries (*27*). Our cohort, which included participants from five regions across two continents, revealed marked site-specific variation in both microbiome composition and MAIT cell frequency and phenotype. The lower frequency of circulating MAIT cells observed in African donors was associated with a more activated phenotype, raising the possibility of tissue redistribution, activation-associated cell loss or altered homeostatic regulation. These possibilities remain to be tested directly. In contrast to a previous study (*31*), we observed a higher abundance of circulating MAIT cells in males than females, suggesting that previous observations made in Western cohorts may not be generalisable across diverse populations.

In contrast to most T cell antigens and due to their small size, 5-OP-RU and its precursor 5-A-RU can diffuse from the intestinal lumen into the circulation and influence MAIT cells at distant sites (*7, 8, 25*). These properties justified quantification of MAIT antigens in faecal samples. Existing approaches are either insufficiently sensitive for this application or challenging to apply to non-sterile, PAMP-rich biological matrices. We therefore developed a cell-based assay using THP-1 cells overexpressing human MR1 together with a soluble recombinant high-affinity MAIT TCR. This strategy takes advantage of the preferential surface expression of antigen-loaded MR1 (*32*) and adds a second layer of selectivity through recognition of MR1-ligand complexes by the engineered MAIT TCR, enabling antigen detection in the picomolar range and thereby revealing novel biology. The association between faecal antigen concentration and circulating MAIT cell phenotype supports a model in which soluble microbial metabolites contribute to systemic MAIT cell tuning.

The strongest microbial associations with faecal MAIT antigen concentration were linked to abundance of the riboflavin biosynthesis pathway in archaea rather than bacteria or fungi. This was unexpected, as 5-OP-RU was initially discovered in bacteria (*5, 6*) and has since been studied largely in that context. The absence of archaea from the MAIT cell literature may reflect the broader underrepresentation of the human archaeome in host–microbiome research, partly because no archaeal pathogens have been clearly identified and because archaeal detection, culture and annotation remain technically challenging (*1, 33*). Methanogenic archaea, including the common human commensal *Methanobrevibacter smithii*, are keystone members of the gut ecosystem, consuming hydrogen generated by bacterial fermentation and thereby supporting syntrophic microbial metabolism (*1*). Among the archaeal species detected in our cohort, *M. smithii* was most abundant and prevalent. Notably, *M. smithii* and related species possess a metabolic pathway capable of generating 5-A-RU as an intermediate in the biosynthesis of factor 420, a coenzyme required for hydrogenotrophic methanogenesis (*34, 35*). The presence of this pathway suggested that archaea might activate MAIT cells through MR1, either by releasing antigenic metabolites into the extracellular environment or following uptake by antigen-presenting cells. Using our novel assay, we found that *M. smithii* culture supernatants contained 5-A-RU/5-OP-RU at levels comparable to or higher than those measured for bacterial commensals and pathogens in our panel. Closely related *Methanobrevibacter* and *Methanobacterium* species sharing this pathway also produced detectable antigenic activity, although at lower levels. MAIT cell activation was further observed after incubation of fixed microbes with autologous antigen-presenting cells, and this response was abolished by MR1 blockade.

The quality of MAIT cell activation induced by *M. smithii* differed from that induced by bacteria. Whereas *M. smithii* promoted clear upregulation of activation markers, cytokine production was comparatively limited. This suggested that archaeal stimulation may resemble isolated TCR triggering more closely than the combined TCR and innate cytokine stimulation typically induced by bacteria. This profile is consistent with the limited capacity of archaea, lacking key bacterial and fungal PAMPs, to engage canonical inflammatory pattern-recognition pathways in antigen-presenting cells (*1, 36, 37*). In line with this model, myeloid-derived MAIT-stimulating cytokines were strongly induced in the *E. coli* condition but not following stimulation with *M. smithii*.

Together, these findings provide evidence that an abundant human T cell population can sense the archaeal commensal *M. smithii*. The mechanisms by which archaeal metabolites are released, captured and presented by MR1 remain to be defined. Future studies should determine the contribution of antigen-processing pathways and innate receptors to MAIT cell responses against archaea. Based on the activation profile induced by *M. smithii*, we propose that repeated TCR stimulation in the relative absence of inflammatory cytokines may contribute to a distinct MAIT cell state, reflected in the individuals in our African cohorts. Rather than promoting a classical TH1-like inflammatory program, chronic exposure to archaeal or microbiota-derived MR1 ligands may favour functions linked to tissue homeostasis and repair. Such functions have been described previously for MAIT cells and appear to be strongly influenced by TCR-dependent signals (*9, 22*). More broadly, our findings identify archaeal metabolism as an unappreciated source of MAIT cell antigens and suggest that the human microbiome can shape systemic T cell immunity through diffusible metabolites produced by non-bacterial members of the intestinal ecosystem.

Future studies will be needed to determine how microbiota-derived MR1 ligands alter MAIT cell behaviour and how this contributes to their functions in health and disease. MAIT cells have been shown to exert protective effects during bacterial (*15*–*21*) and viral (*13*) infection and to contribute to vaccine responses (*38*), but they have also been implicated in the pathogenesis of autoimmune and inflammatory diseases (*39, 40*). Our findings raise the possibility that diffusion of microbiome-derived MR1 ligands maintains MAIT cells in a low-inflammatory state, potentially favouring tissue repair over pathogenic inflammation. Antigen production varies across microbiome configurations, contributing to the wide inter-individual variability in MAIT cell abundance and phenotype and potentially pointing to a mechanistic link between changes in human microbiomes and broader immune homeostasis across populations.

## General

We would like to acknowledge Carole Perot and Gloria Mesa Gil at Immunocore for the initial MAIT TCR affinity enhancement; Daniel Fonseca at Immunocore for discussion on the MAIT TCR; Jonathan Webber at the flow cytometry facility at the Kennedy Institute of Rheumatology for support with high-dimensional flow cytometry panel design and optimisation; Helen Ferry at the Flow Cytometry Facility, Experimental Medicine Division, University of Oxford for support with the flow cytometric analysis; Dini Senanayake for high-performance computing cluster support; the Oxford Vaccine Centre Biobank; The Translational Gastroenterology and Liver Unit Biobank; Nicola Borthwick at the Jenner Institute for assistance in accessing samples; Emile Pernet, Julia Dabrowka Podolan and Tabea Oswald for their contributions during their internships; the NIH Tetramer Core Facility (contract number 75N93020D00005) for providing tetramers, and the volunteers who donated the samples used in this study.

## Author contributions

Conceptualization: NF-P, MJL, NP, PK, JSJ

Formal analysis: NF-P

Funding acquisition: PK, JSJ

Investigation: NF-P, MJL, IWDM, CJSP

Methodology: NF-P, MJL, CJSP, MS, RJS, RS, KB

Project administration: WJ, FK, WK, EJS, PC, TH

Resources: BS

Supervision: MCC, CFP, NM-P, PK, JSJ

Visualisation: NF-P, MJL, IWDM

Writing – original draft: NF-P, MJL

Writing – review & editing: NF-P, MJL, IWDM, CJSP, MS, RJS, RS, KB, BS, WJ, FK, WK, EJS, MCC, CFP PC, TH, NMP, PK, JSJ

## Data availability

Raw spectral flow cytometry phenotyping data and the associated metadata have been deposited in ImmPort under accession Johnson_kennedy.ox. Shotgun metagenomic sequencing data generated in this study have been deposited in the European Nucleotide Archive (ENA) under the project accession number PRJEB124982. Requests for materials should be addressed to Paul Klenerman or Jethro S. Johnson.

