## Supplemental Figures and Tables for "Gut archaea drive human MAIT cell responses"

**The PDF file includes:**

Materials and Methods
Figs. S1 to S8
Tables S1 to S2

### Materials and Methods

#### *Ethical approval*

Cryopreserved faeces and peripheral blood mononuclear cells (PBMCs) from healthy donors from Oxford, UK (n = 20 donors for faeces; subset of n = 10 donors for PBMCs); Lusaka, Zambia (n = 22); Nairobi, Kenya (n = 22); Kilifi, Kenya (n = 22) and Masaka, Uganda (n = 22) were accessed through the Oxford Vaccine Centre (OVC) Biobank. Participants gave their informed consent for their samples to be stored in the OVC Biobank. The Biobank has ethical approval (21/SC/0161) to release samples to researchers at Oxford University for projects meeting the objectives of the Biobank without further ethical review. Faecal samples and PBMCs from UK donors (n = 10) were accessed from the trial HIV-CORE 005.2 (NCT04586673). The trial was granted a favourable opinion by the East of England–Cambridge Central Research Ethics Committee (Ref:20/EE/0036) and approved by the UK Medicines and Healthcare Products Regulatory Agency (CTA 21584/0425/001-0001; EudraCT 2019-003973-25; IRAS 263882). Further faecal samples from UK donors (n = 10) were accessed from the trial HIV-CORE 005.1 (NCT04563377). This trial was granted a favourable opinion by the East of England–Cambridge Central Research Ethics Committee (Ref:19/EE/0323) and approved by the UK Medicines and Healthcare Products Regulatory Agency (EudraCT 2019-000621-47; IRAS 261129). Faecal samples and PBMCs from African donors were accessed from the trial HIV-CORE 006 (NCT04553016; PACTR202006495409011). The arm of the study performed at the Center for Family Health Research Zambia, Lusaka was approved by the University of Zambia Biomedical Research Ethics Committee (UNZABREC 495-2019) and Zambia Medicines Regulatory Authority (CT 098), the National Biosafety Authority (NBA/101/16/1), the National Health Research Authority and by Oxford Tropical Research Ethics Committee (OXTREC 56-19). The arm of the study performed at MRC/UVRI and London School of Hygiene and Tropical Medicine Uganda Research Unit, Masaka was approved by Uganda Virus Research Institute REC (GC/127/20/06/761), Uganda National Council for Science and Technology (HS844ES), and National Drug Authority (CTC 0171/2021). The arm performed at Kenya Medical Research Institute-Wellcome Trust Research Programme Kilifi (KWTRP) was approved by Clinical Science Committee (CSC 180), KEMRI Scientific and Ethics Review Unit (SERU 4025), National Commission for Science, Technology & Innovation (NACOSTI/P/23/23155/) and Pharmacy and Poisons Board (PPB/ECCT/20/10/01/2021). The arm performed at KAVI-Institute for Clinical Research (KAVI-ICR), University of Nairobi, Nairobi was approved by Kenyatta National Hospital-University of Nairobi Ethics Research Committee (P863/10/2019), NACOSTI (NACOSTI/286477) and PPB (PPB/ECCT/20/06/09/2020). All studies were conducted according to the principles of the Declaration of Helsinki (2008) and complied with the International Conference on Harmonization Good Clinical Practice guidelines and Good Participatory Practices. PBMCs for assays in Figure 4 were accessed from the Translational Gastroenterology Unit (TGLU) Biobank in Oxford, UK (NHS REC 21/YH/0206) with patient consent.

#### *Flow cytometry profiling of human PBMCs*

##### Isolation, cryopreservation and thawing PBMCs

Blood was drawn into heparinized vacutainers (Becton Dickinson, NJ, USA) and PBMCs isolated within 6 hours. PBMCs were frozen overnight at -80°C using Nalgene® Mr Frosty containers (Sigma-Aldrich, MA, USA) then transferred to liquid nitrogen vapour within 48 hours for long-term storage. PBMCs in cryovials were thawed in a water bath at 37°C then added dropwise to warmed R10 medium (RPMI 1640 medium (Thermo Fisher Scientific, MA, USA) supplemented with 10% FBS (R&D Systems, MN, USA)) and penicillin-streptomycin (Life Technologies, CA, USA). Cells were washed in R10, counted and resuspended to 20 million live cells/mL. 100 µL cells were plated in a V-bottomed 96-well plate (Corning, ME, USA) for immediate phenotypic staining or stimulated as below.

**Stimulation of PBMCs with anti-CD3 and anti-CD28 or IL-12 and IL-18**
For TCR stimulation, 96-well flat-bottom MaxiSorp Nunc-Immuno plates (Thermo Fisher Scientific, MA, USA) were coated with 100 µL purified anti-human CD3 antibody (clone OKT3, BioLegend, CA, USA) per well, prepared in PBS (Thermo Fisher Scientific, MA, USA) to a concentration of 2.5 µg/mL and incubated overnight at 4°C. The anti-CD3 antibody was discarded and the plate washed twice with PBS. Wells were blocked by incubating with 200 µL R10 for two hours at 37°C. The R10 was discarded and 100 µL freshly thawed cells plated per well with purified anti-human CD28 antibody (clone CD28.2, BioLegend, CA, USA) prepared in R10, to give a final concentration of 1 µg/mL anti-CD28 per well. For cytokine stimulation, 100 µL freshly thawed cells were plated per well of a MaxiSorp plate with 100 µL stimulation cocktail containing 0.1 µg/mL recombinant human interleukin 18 (IL-18) (R&D Systems, MN, USA) and 0.1 µg/mL recombinant human interleukin 12 (IL-12) (R&D Systems, MN, USA) in R10. For unstimulated controls, 100 µL R10 was added in place of the stimulation cocktail. Cells were incubated at 37°C for 24 hours. Four hours before the end of the incubation period, eBioscience™ Brefeldin A Solution (1,000X) (Thermo Fisher Scientific, MA, USA) was added to give a final dilution of 1:1,000. Cells were transferred to a V-bottomed 96-well plate for staining.

**Staining for flow cytometry**
Cells were centrifuged at 2,200 rpm for 2 minutes, resuspended in 200 µL FACS buffer (PBS supplemented with 0.1% bovine serum albumin and 5 mM UltraPure™ EDTA (Thermo Fisher Scientific, MA, USA)) to wash, then centrifuged again. Cells were then incubated with Human TruStain FcX™ (BioLegend, CA, USA) at a dilution of 1:400 and LIVE/DEAD™ Fixable Blue Stain (Thermo Fisher Scientific, MA, USA) at a dilution of 1:700 in a total volume of 100 µL PBS for 30 minutes at 4°C, protected from light. Cells were washed twice in FACS buffer then incubated for 30 minutes with the antibody cocktail for surface staining, prepared to a total volume of 50 µL in Brilliant Stain Buffer (BD Biosciences, CA, USA). Antibodies for the entire panel are listed in Table S2. After surface staining, cells were washed twice then incubated for 20 minutes in Cytofix/Cytoperm™ Fixation and Permeabilization Solution (BD Biosciences, CA, USA) at 4°C, protected from light. Cells not receiving a 24-hour stimulation were washed twice with FACS buffer and resuspended in 200 µL FACS buffer ready for acquisition. Stimulated cells were washed twice with Perm/Wash™ Buffer (BD Biosciences, CA, USA) then incubated in the antibody cocktail for intracellular cytokine staining, prepared to a total volume of 50 µL in Perm/Wash buffer, for 30 minutes at 4°C, protected from light. Cells were washed twice with Perm/Wash buffer then resuspended in FACS buffer for acquisition. Both cell-based and bead-based single-colour controls were prepared with every batch of samples. For cell-based controls, 250,000 cells were plated per well. Cells used for single-colour controls for activation markers and cytokines were stimulated for 24 hours with anti-CD3 and anti-CD28 as above, whilst cells for phenotypic markers were plated for 24 hours in R10. For bead-based single-colour controls, one drop of UltraComp eBeads™ Compensation Beads (Thermo Fisher Scientific, MA, USA) was added per well. Cell-based and bead controls were stained in parallel to cells stained with the full staining

cocktail, in the same concentrations as antibodies in the full cocktail. An unstimulated unstained control and an unstained control stimulated with anti-CD3 and anti-CD28, both containing one million cells, were prepared for autofluorescence extraction. PBMCs from the same blood cone were stained with the full panel for every stimulation condition in every randomised batch of 15-20 samples for batch correction.

**Flow cytometry data acquisition**

Data were acquired on a standard 5-laser 16UV/16V/14B/10YG/8R Aurora spectral flow cytometer (Cytek Biosciences, CA, USA). Data were acquired and spectrally unmixed with autofluorescence extraction using SpectroFlo software (v3.3.0, Cytek Biosciences, CA, USA).

**Quality control and gating of spectral flow cytometry data**

Data were prepared for high dimensional analysis following a workflow described by den Braanker *et al.* (*41*). Unmixed FCS files were manually gated in FlowJo software (v10.10.0, BD Biosciences, CA, USA), according to the gating strategy in fig. S1. Firstly, a time gate was used to exclude noise due to inconsistencies in flow rate, then singlets and $CD45^+$ live cells were selected. The population was then gated for lymphocytes and $CD19^+$ and $CD14^+$ cells were excluded. The exported FCS 3.1 files were imported into OMIQ software (Dotmatics, MA, USA). OMIQ was used to create workflows incorporating the following tools, with a separate workflow for *ex vivo* stained cells to stimulated cells. Firstly, data underwent *arcsinh* transformation with a fixed cofactor of 6,000 for all features. Automatic quality control was performed using PeacoQC (v1.14.0), configured with the default parameters, to remove anomalies or outlier events not removed by manual gating. Batch correction was then performed. For the stimulated cells workflow, the unstimulated and stimulated controls were concatenated within each batch. Then, elbow metaclustering was performed with FlowSOM (v3.21) using default parameters (xdim = 10, ydim = 10, number of training iterations = 10, distance metric = Euclidian) to cluster control samples across batches. These clusters informed batch effect removal using CytoNorm (v2.0.9) using default parameters (number of quantiles = 101). Following quality control, samples were gated for major cell populations using the gating strategy outlined in fig. S1. Cells which received 24-hour stimulation were manually gated for MAIT cells, then gated for phenotypic markers and cytokines according to the gating strategy in fig. S2. The abundance of MAIT cells positive for each marker in the unstimulated condition was subtracted from each stimulated condition, for each donor.

**Flow cytometry data visualisation and statistical analysis**

Elbow metaclustering was performed on MAIT cells based on expression of CD4, CD8, CD25, PD-1, CD47, CD38, HLA-DR, CD45RA, CCR7 and CD69 using FlowSOM. The FlowSOM default parameters were used (xdim = 10, ydim = 10, number of training iterations = 10, distance metric = Euclidian). Expression of markers by each of the 16 clusters was visualised by a heatmap of min-max scaled median fluorescence intensity of the markers. Clusters were manually grouped together by shared expression patterns of activation markers. Two-dimensional flow cytometry plots for gating strategies were generated in FlowJo and OMIQ. All other visualisations, and all statistical analyses, were generated in R (v4.3.1).

*Faecal microbiome analysis*

**Faecal sample collection and storage**

Participants self-collected a pea-sized stool sample using a collection kit. For the HIV-CORE 005.1 trial the collection kit consisted of a paper carrier bag, a pair of disposable gloves, a thermal lined transport box (Thermal Control Unit TC-565, Air Sea Containers, Merseyside, UK), two ThermoSafe PolarPack ice packs (Sonoco ThermoSafe, IL, USA), a Faeces Catcher (Zymo Research, CA, USA) faeces collection device, a faeces tube (Sarstedt, Nümbrecht, Germany) and a BioBag transporter (Air Sea Containers, Merseyside, UK). For the HIV-CORE 005.2 and HIV-CORE 006 trials, participants were provided with a stool collection bag containing a stool pot, a transport container with absorbent tissue, a Fe-Col® Faeces Collection Device (Alpha Laboratories, Hampshire, UK) and gloves. Faecal samples were stored at 2-8°C and returned to the clinic within 24 hours. Samples were transported to the lab from the clinic in cool bags with a cool pack. Upon receipt at the lab, faecal samples were divided into four labelled aliquots containing approximately 300 mg in a Class II safety cabinet following aseptic technique. Aliquots were transferred immediately to a -80°C freezer for storage. Samples were shipped on dry ice, avoiding thawing during shipment.

**Measurement of microbial load by flow cytometry**

Faecal samples were thawed at room temperature and transferred into pre-weighed O-ring tubes using a sterile pipette tip and weighed to calculate faecal mass. 1 mL sterile PBS was added, and faeces manually mixed using a sterile spreader. The tube was vortexed at maximum speed on a Vortex Adapter for 24 tubes (QIAGEN, Venlo, Netherlands) coupled to a Vortex-Genie 2 (QIAGEN, Venlo, Netherlands) for 10 minutes, until the faeces were completely homogenised. The sample was centrifuged at 300 × *g* for 5 minutes at 4°C to pellet large debris, leaving microbes in suspension. The supernatant was filtered through a 70 µM cell strainer then centrifuged to pellet microbial cells at 16,000 × *g* for 5 minutes at 4°C. The pellet was resuspended in 1 mL sterile PBS by pipetting, to wash. The centrifugation and wash steps were repeated for a total of 4 washes. The sample was then serially diluted to a final concentration of 1:20 000 and stained with SYTO9 and propidium iodide from the LIVE/DEAD BacLight Bacterial Viability Kit for Microscopy and Quantitative Assays (Thermo Fisher Scientific, MA, USA) according to the manufacturer's instructions. Stained samples were incubated or 15 minutes at room temperature, protected from light, then 100 µL was acquired on a standard 5-laser 16UV/16V/14B/10YG/8R Aurora spectral flow cytometer (Cytek Biosciences, CA, USA) within 15-30 minutes of adding the stain, with a threshold of 3,000 in the B1 detector. Total gated events were multiplied by 200,000 then normalised to count/mg stool.

**Faecal DNA extraction**

Faecal DNA was extracted from faeces thawed at room temperature using the ZymoBIOMICS DNA Miniprep Kit (Zymo Research, CA, USA), according to manufacturer's instructions. For a subset of African samples, DNA yield was poor following two attempts at extraction. For these samples, DNA was extracted from a different faecal aliquot using the QIAamp PowerFecal Pro DNA Kit (QIAGEN, Venlo, Netherlands) according to manufacturer's instructions. Additional samples with higher yield were also re-extracted using the QIAGEN kit to ensure an equal number of samples were extracted with each kit from each study site. Sample homogenisation was performed for all samples using a Vortex Adapter for 24 tubes (QIAGEN, Venlo, Netherlands) coupled to a Vortex-Genie 2 (QIAGEN, Venlo, Netherlands) for 20 minutes at maximum speed.

DNA A260/A280 and A260/A230 ratios were measured on a NanoDrop Microvolume Spectrophotometer (Thermo Fisher Scientific, MA, USA) and DNA quantification was performed on a Qubit Fluorometer using the Qubit™ dsDNA HS Assay Kit (Thermo Fisher Scientific, MA, USA). DNA was diluted to a final concentration of 20-40 ng/µL in elution buffer before sequencing. Each randomised batch of 12-24 samples included one blank negative control. 2-3 positive controls were prepared for each extraction kit in parallel with sample extractions, using the ZymoBIOMICS Microbial Community Standard (Zymo Research, CA, USA).

**DNA library preparation and shotgun metagenomic sequencing**

DNA library preparation and shotgun metagenomic sequencing were performed at Oxford Genomics Centre for UK samples. Material was quantified using Qubit (Invitrogen, CA, USA) or PicoGreen (Thermo Fisher, CA, USA) on the FLUOstar OPTIMA plate reader (BMG Labtech, Ortenberg, Germany) and normalised to 100 ng. Fragmentation was performed by mechanical shearing to an average size of 350 bp using an EpiSonic MultiFunctional Bioprocessor (Epigentek, NY, USA; amplitude 40, process time 3 min 20 sec, pulse on/off 20 sec). Library preparation was performed using the NEBNext Ultra DNA library prep kit for Illumina (New England Biolabs, MA, USA) and standard Illumina multiplexing adapters with minor modifications to manufacturer's protocol. Libraries were PCR amplified (12 cycles) on a Tetrad (Bio-Rad, CA, USA) using in-house unique dual indexing primers, based on Lamble *et al* (*42*). Post-PCR purification performed using Agencourt Ampure XP (Beckman Coulter, CA, USA; ratio 1:0.75). Individual libraries were normalised using PicoGreen (Thermo Fisher, CA, USA) or Quantifluor (Promega, WI, USA) and pooled together accordingly. The size profile of the pooled library was analysed on the 2200 or 4200 TapeStation (Agilent, CA, USA). The pooled library was quantified using Qubit (Invitrogen, CA, USA) and diluted to ~10 nM for storage. The 10 nM library was denatured and further diluted prior to loading on the sequencer. Paired end sequencing was performed using a NovaSeq 6000 150 bp platform (Illumina, CA, USA), NovaSeq 6000 S2 Reagent Kit (300 cycles) or NovaSeq 6000 S4 Reagent Kit (300 cycles)). The closure of Oxford Genomics Centre prevented the sequencing of samples from African volunteers following an identical protocol to the UK samples. Instead, a commercial sequencing provider (Novogene UK, Cambridge, UK) performed library preparation using the Novogene NGS DNA Library Prep Set and shotgun metagenomic sequencing on a NovaSeq X Plus instrument (Illumina, CA, USA) with 150-bp paired-end reads.

**Quality control and pre-processing of shotgun metagenomic sequencing data**

Pipelines developed by the Oxford Centre for Microbiome Studies (OCMS) (*43*) were used for bioinformatic processing of shotgun sequencing data. Quality of raw sequencing data was assessed using FastQC (v0.11.9) and MultiQC (v1.9), implemented in OCMS pipeline_fastqc.py. Pre-processing of reads was then performed using OCMS pipeline_preprocess.py, based on the original Human Microbiome Project protocol (*44*). First, identical PCR duplicates were removed using the cd-hit-dup program from CD-HIT (v4.8.1). Next, adapter contamination was removed using Trimmomatic (v0.39) with the settings ILLUMINACLIP:"adapters/custom/novogene.fa":3:30:10:8:True LEADING:25 TRAILING:25 MINLEN:36. Host contamination was then removed using the Best Match Tagger (BMTagger) (v3.101) and finally, low complexity regions were masked using BBMap (v38.90).

**Taxonomic assignment and functional profiling**

Taxonomic assignment and functional profiling of pre-processed fastq files was performed using OCMS pipeline_humann3.py. Pre-processed reads were annotated using HUMAnN 3.8, based on DIAMOND (v0.9.36) and Bowtie2 (v2.4.1), with the ChocoPhlAn database (mpa_v31_CHOCOPhlAn_201901) and UniRef90 (v201901b) protein database to quantify abundance of metabolic pathways. The annotation results were mapped to MetaCyc functional pathways (*30*) using the "humann3_regroup_table" script. Reads aligning to MetaCyc pathways were normalized to sequencing coverage and reported as copies per million reads (cpm). MetaPhlAn (v3.1) was used to classify reads to bacterial species.

**Prevalence-abundance filtering**

Phylum-, genus- and species-level taxonomic abundance tables were filtered to keep taxa with a prevalence $\geq$ 5% or a relative abundance $\geq$ 0.1% in at least one sample. To prevent disproportionate removal of taxa found exclusively in UK microbiomes (given that the dataset largely consisted of African microbiomes), prevalence-abundance filtering was initially performed separately for each study site. Only taxa identified for removal from every study site were filtered from the whole dataset, resulting in 15 phyla, 169 genera and 595 species. Relative abundances were then re-scaled so the total relative abundance for each sample was equal to 100%. MetaCyc pathways were filtered across the whole dataset at a prevalence threshold of 5%, leaving 476 pathways.

**Analysis of microbiome diversity**

Species richness were calculated using the specnumber() function from the *vegan* R package (v2.6.10). To assess beta diversity, Bray-Curtis dissimilarity (BCD) was calculated on genus-level relative abundance and MetaCyc pathway abundance using the vegdist() function from the *vegan* R package (v2.6.10). A Principal Coordinate Analysis (PCoA) was performed on each BCD matrix using the base R cmdscale() function with k=10. Coordinates from these 10 axes were extracted for plotting and statistical analysis.

**Association between microbiome features and MAIT cell features**

Linear mixed-effects models were used to assess the relationship between MAIT cell abundance or activation and microbial and demographic predictors. Models were fitted using the lmer() function from the *lme4* R package (v1.1.37) with significance of predictors assessed by type III F-tests using the *lmerTest* R package (v3.1.3). To calculate the variance in a particular immune phenotype explained by gut microbiome composition or MetaCyc pathway abundances, a linear model was fit using the base R function lm() on the first 10 PCoA axes generated from a Bray-Curtis dissimilarity matrix. R-squared and a p-value were extracted for each model using the summary() function from the package *stats4* (v4.3.1). To explore associations between all 476 MetaCyc pathway abundances and each MAIT cell phenotypic marker, Spearman's rank correlation coefficient ($\rho$) was calculated using the cor.test() function from the R package *stats* (v4.3.1) with the Benjamini-Hochberg correction. Correlations were visualised using the Heatmap() function from the package ComplexHeatmap (v2.18.0).

*Quantification of MAIT cell antigens*

**Production of high affinity MAIT TCR**

The high affinity MAIT TCR (MR1-01-S1-a10b10) was fused to a rabbit Fc domain, with each TCR chain C-terminally fused to one half of the Fc domain (knob-in-hole). Proteins were expressed following transient transfection of pCDNA3.1 constructs into the ExpiCHO Expression System (ThermoFisher Scientific) and purified using Protein A-affinity (MAbSelect SuReTM (Cytiva, Marlborough, MA, USA)) and size exclusion chromatography (S200 10/300 column) (in Dulbecco's PBS).

**Generation of an MR1-overexpressing THP-1 cell line**

THP-1 cells were maintained in RPMI 1640 medium supplemented with 10% heat-inactivated fetal bovine serum, 2 mmol/L L-glutamine (all Gibco, Thermo Fisher Scientific, MA, USA), 100 U/mL penicillin, and 100 µg/mL streptomycin (Sigma-Aldrich, MO, USA). Cells were cultured at 37°C in a humidified atmosphere containing 5% $CO_2$ and maintained at a density between $10^5$ and $3 \times 10^6$ cells/mL. To generate a stable MR1-overexpressing cell line, THP-1 cells were transduced with lentiviral vectors encoding human MR1 isoform 1 (NP_001372090.1). Three sequential rounds of lentiviral transduction were performed to maximise the proportion of MR1-expressing cells. For each round, $3 \times 10^6$ THP-1 cells were incubated with the MR1-encoding lentiviral preparation in the presence of 8 µg/mL polybrene (Sigma-Aldrich, MA, USA). Lentiviral transduction was enhanced by spinoculation at 1,100 × *g* for two consecutive 45-min periods at 33°C. Following the second round of transduction, cells were selected with puromycin (Gibco, Thermo Fisher Scientific, MA, USA) at a final concentration of 0.8 µg/mL. After the third and final round of transduction, cells were allowed to recover for 48 h before selection with blasticidin (Gibco, Thermo Fisher Scientific, MA, USA) at a final concentration of 8 µg/mL. Blasticidin selection was continued for 7 days, with regular replacement of the selection medium, until the non-transduced control cells were no longer viable. Cell clones were then generated by limiting dilution. The resulting stable cell population, hereafter referred to as THP-1–MR1 cells, was expanded and cryopreserved. Cell-surface MR1 expression was confirmed by flow cytometry using an anti-human MR1 monoclonal antibody (clone 26.5).

**Preparation of 5-OP-RU standards**

The MR1 ligand 5-(2-oxopropylideneamino)-6-D-ribitylaminouracil (5-OP-RU) was generated immediately before use by reacting 5-amino-6-D-ribitylaminouracil (5-A-RU; cat. no. 39898; Cayman Chemical, MA, USA) with methylglyoxal (cat. no. M0252; Sigma-Aldrich, MA, USA). Briefly, 1 µL of a 4% (560 mmol/L) methylglyoxal solution prepared in water was added to a 3 mmol/L solution of 5-A-RU prepared in dimethyl sulfoxide (DMSO; Sigma-Aldrich, MA, USA). The reaction mixture was incubated on ice for 4 minutes and used immediately without further purification. The nominal concentration of the resulting 5-OP-RU stock was calculated from the amount of 5-A-RU present in the reaction mixture, assuming quantitative conversion of 5-A-RU to 5-OP-RU. Serial dilutions were prepared immediately before each experiment in assay medium to generate the 5-OP-RU standard curve.

**Faecal sample lysis**

Fecal samples were transferred into tubes containing 0.1 mm zirconium beads (Sigma-Aldrich, MA, USA) and weighed. Samples were diluted 1:3 in sterile DPBS (Gibco, Thermo Fisher

Scientific, MA, USA) (mass to volume) and homogenised using a FastPrep-24 instrument for 3 cycles of 20 seconds at 6 m/s. Tubes were kept on ice before lysis and immediately after lysis. Following homogenisation, samples were centrifuged at 17,000 × *g* for 10 minutes at 4°C. The supernatant was then collected for downstream analysis. Faeces from *Blautia coccoides* (a riboflavin-incompetent species) mono-colonised mice were used as a negative control for assay background removal.

**MR1 ligand-loading assay**

The assay medium consisted of 80% dialysed foetal calf serum and 20% RPMI 1640 medium (v/v), containing 0.1 × pyridoxal and 0.1 × folate. THP-1–MR1 cells were harvested, washed once, and resuspended in assay medium at a concentration of $10^7$ cells/mL. A total of $10^5$ cells, corresponding to 10 µL of cell suspension, was dispensed into each well of a 96-well U-bottom plate. Biological samples were diluted 1:20 in assay medium before addition to the cells. In parallel, freshly prepared 5-OP-RU standards were serially diluted in the same assay medium. THP-1–MR1 cells were incubated with diluted biological samples, 5-OP-RU standards, or vehicle controls for 3.5 h at 37°C in a humidified atmosphere containing 5% $CO_2$ to allow ligand uptake and loading onto cell-surface MR1.

**Soluble MAIT TCR staining and Fab-fragment detection**

Following antigen loading, cells were washed once and incubated for 30 min at 37°C in blocking buffer containing 20% foetal calf serum and Fc-blocking reagent diluted 1:100. Cells were subsequently stained with the soluble high affinity MAIT TCR MR1-01-S1-a10b10-rbFc produced at Immunocore (Oxfordshire, UK) at final staining concentration of approximately 2.0 nmol/L. Cells were incubated with the soluble TCR for 30 min at 37°C and subsequently washed to remove unbound reagent. Cell-bound soluble TCR was detected using two distinct fluorescently labelled Fab fragments recognising the soluble rabbit-Fc part of the TCR detection construct. Donkey anti-Rabbit phycoerythrin (cat no. 711-116-152, JIR, Cambridge, UK) was diluted 1:250, and Goat anti-Rabbit Coralite 647 (cat no. SA00014-9, Proteintech, IL, USA) was diluted 1:300. Secondary staining was incubated for 20 min at 21°C, protected from light. Following staining, cells were washed and resuspended in 100 µL of flow-cytometry buffer consisting of PBS supplemented with 2% foetal calf serum, 0.1% sodium azide (Sigma-Aldrich, MA, USA), 0.5 mM EDTA (Sigma-Aldrich, MA, USA), and 0.01% Tween 20 (Sigma-Aldrich, MA, USA). The gating strategy and an example of a standard curve can be seen in fig. S7, A and C.

**Spectral flow cytometric acquisition and data analysis**

Samples were acquired using a 5-laser (16UV/16V/14B/10YG/8R) Aurora spectral flow cytometer (Cytek Biosciences, CA, USA). A consistent gating strategy was applied to all standards and biological samples. Cellular debris was excluded based on forward- and side-scatter characteristics, and doublets were excluded using forward-scatter area and height parameters. The median fluorescence intensity (MFI) of each Fab-fragment signal was determined within the final THP-1–MR1 cell population on the diagonal scale for parameters "YG1"-"R1" and "B4"-"R3". The assay signal was calculated as the sum of the Geo-MFIs measured in the PE and Coralite 647 detection channels using R to sum parameter "B4-A", "B5-A", "YG1-A", "YG2-A", "YG3-A", "YG4-A", "YG5-A", "YG6-A", "YG7-A", "YG8-A", "YG9-A", "R1-A","R2-A","R3-A","R4-A","R5-A","R6-A","R7-A" and "R8-A". The final values were interpolated from a standard curve fitted using a four-parameter logistic (4PL) regression to determine the initial analyte concentration, which was subsequently normalised to stool weight and expressed per mg of stool.

#### *Culture of bacteria and archaea*

##### Bacterial culture

*Akkermansia muciniphila* and *Clostridium innocuum* were cultured in a COY anaerobic chamber (Coy Laboratory Products, MI, USA) (gas cylinder composition 0% $O_2$, 10% $CO_2$, 10% $H_2$, 80% $N_2$; chamber gas mix 150 ppm $O_2$ and 2.5% $H_2$) using plasticware de-gassed for a minimum of 12 hours. Strains were grown in 5 mL AF medium as previously described (*45*) in 13 mL vented cap tubes (Sarstedt, Nümbrecht, Germany) for 16 h. Bacteria were counted by staining with SYTO9 and propidium iodide and acquiring by flow cytometry as described above. Archaeal and bacterial cultures were centrifuged at 16,000 × *g* for 5 minutes at 4°C. Supernatants and pellets were stored at -70°C for downstream assays. All other bacterial strains were cultured in tryptone soya broth (Oxoid, Basingstoke, UK) supplemented with riboflavin (CHEMCRUZ, Dallas, TX, USA) at a final concentration of 20 µg/mL. Cultures were incubated at 37 °C with orbital agitation at 220 rpm. Bacterial growth was monitored by measuring the optical density at 600 nm ($OD_{600}$). Colony-forming unit concentrations (CFU/mL) were estimated from the $OD_{600}$ measurements using an established $OD_{600}$-to-CFU calibration curve.

##### Methanogenic archaeal culture

Archaea were cultured anaerobically in a Whitley A95 Workstation (Don Whitley Scientific, West Yorkshire, UK) maintained at a constant temperature of 39°C and approximately 61% humidity. The chamber atmosphere was sustained using an anaerobic gas mixture (10% $H_2$, 10% $CO_2$ and 80% $N_2$) and $O_2$-free $N_2$ gas for purging cycles. Media and glassware were de-gassed for 24 hours before use. The archaeal species selected were *Methanobrevibacter smithii* (DSM 861 and DSM 2374), *Methanobrevibacter ruminantium* (DSM 1093), *Methanobrevibacter thaueri* (DSM 11995) and *Methanobacterium bryantii* (DSM 863 and DSM 7079). 300 µL active culture was inoculated into 3 mL of DSMZ Methanobacterium Culture Medium 119 and cultured for 72 h in a sealed Hungate tube. Archaeal cells were counted in a fixed volume of liquid culture by flow cytometry. Briefly, an aliquot was washed thrice in sterile reduced PBS, serially diluted 1:100, then acquired on a 3-laser 8R/14B/16V Cytek Northern Lights spectral flow cytometer (Cytek Biosciences, CA, USA). Factor 420-positive and negative archaea were identified based on autofluorescence in the V4 detector and size in the SSC channel.

#### *MAIT cell activation assay using autologous monocytes*

Blood cones were obtained from anonymised healthy blood donors (age and sex not available, used for 36 h incubation experiment) or healthy colleagues (age and sex of donors for 24 h incubation experiment: female [27], female [32] and male [57]; for 36 h incubation experiment: male [35] and female [34]). PBMCs were isolated from whole blood by Ficoll-Paque density-gradient centrifugation (STEM CELL, Cambridge, UK). Cells were resuspended in RPMI medium supplemented with 5% human AB serum and plated overnight to allow monocyte adherence. The following day, adherent cells were separated from the non-adherent fraction and rinsed once with DPBS. Archaea and bacteria were fixed by incubating a washed pellet in 100 µL of 2% formalin (Thermo Fisher Scientific) for 5 minutes at room temperature, followed by two washes with 1 mL DPBS. Adherent cells were then incubated for 2 h at 37 °C with fixed bacteria or archaea at a multiplicity of infection (MOI) of 20. Control cultures were incubated with RPMI alone, whereas antigen-specific stimulation was performed with 2 nM 5-OP-RU. Where indicated, anti-MR1 blocking antibody or isotype control antibody was added at 25 µg/mL after the initial 2 h stimulation. After 1 h, the remaining PBMCs were added back to the adherent cells and co-cultured for 24 h or 36 h at 37 °C. Following incubation, cells were stained with MR1 tetramer-PE for 20 min, followed by surface staining. Cells were then fixed and permeabilised using Cytofix/Cytoperm (BD Biosciences) according to the manufacturer's instructions and stained with the intracellular antibody panel (full antibody panel reported in table S2). Anti-TCR antibodies were included in both the surface and intracellular staining mixtures to account for potential TCR internalisation and to preserve gating consistency. Data were acquired on a 5-laser (16UV/16V/14B/10YG/8R) Aurora spectral flow cytometer (Cytek Biosciences, CA, USA). Dimensionality reduction was performed in R (v4.6.0) using the package flowCore (v2.24.0) on z-scaled median fluorescence intensity of the following markers: MR1 tetramer, CD3, HLA-DR, CD161, CD8, CD279, CD69, CD25, IFN-γ, granzyme B, CD38, CD137 and TNF. Markers expressed on all MAIT cells and used for gating (TCR-Vα7.2) or expressed on very small cell populations (IL-17) were excluded from dimensionality reduction but are shown in fig. S8, C and D. Donor repartition is shown in fig. S8E.

#### Measurement of cytokine concentrations

Supernatants from monocyte-stimulated PBMCs were collected after 24 h and stored at −80°C until cytokine measurement. Cytokine concentrations were determined using the LEGENDplex™ Human Inflammation Panel 1 (13-plex; BioLegend, CA, USA), according to the manufacturer's instructions. Briefly, samples were incubated with fluorescently encoded capture beads for 2 h at room temperature with shaking (800 rpm), followed by incubation with a biotinylated detection antibody cocktail for 1 h at room temperature with shaking. Streptavidin–phycoerythrin (SA-PE) was then added directly to the wells, and samples were incubated for an additional 30 min at room temperature with shaking. Following washing, the beads were analysed by flow cytometry. Cytokine concentrations were determined from the corresponding standard curves using LEGENDplex™ Data Analysis Software (BioLegend, CA, USA).

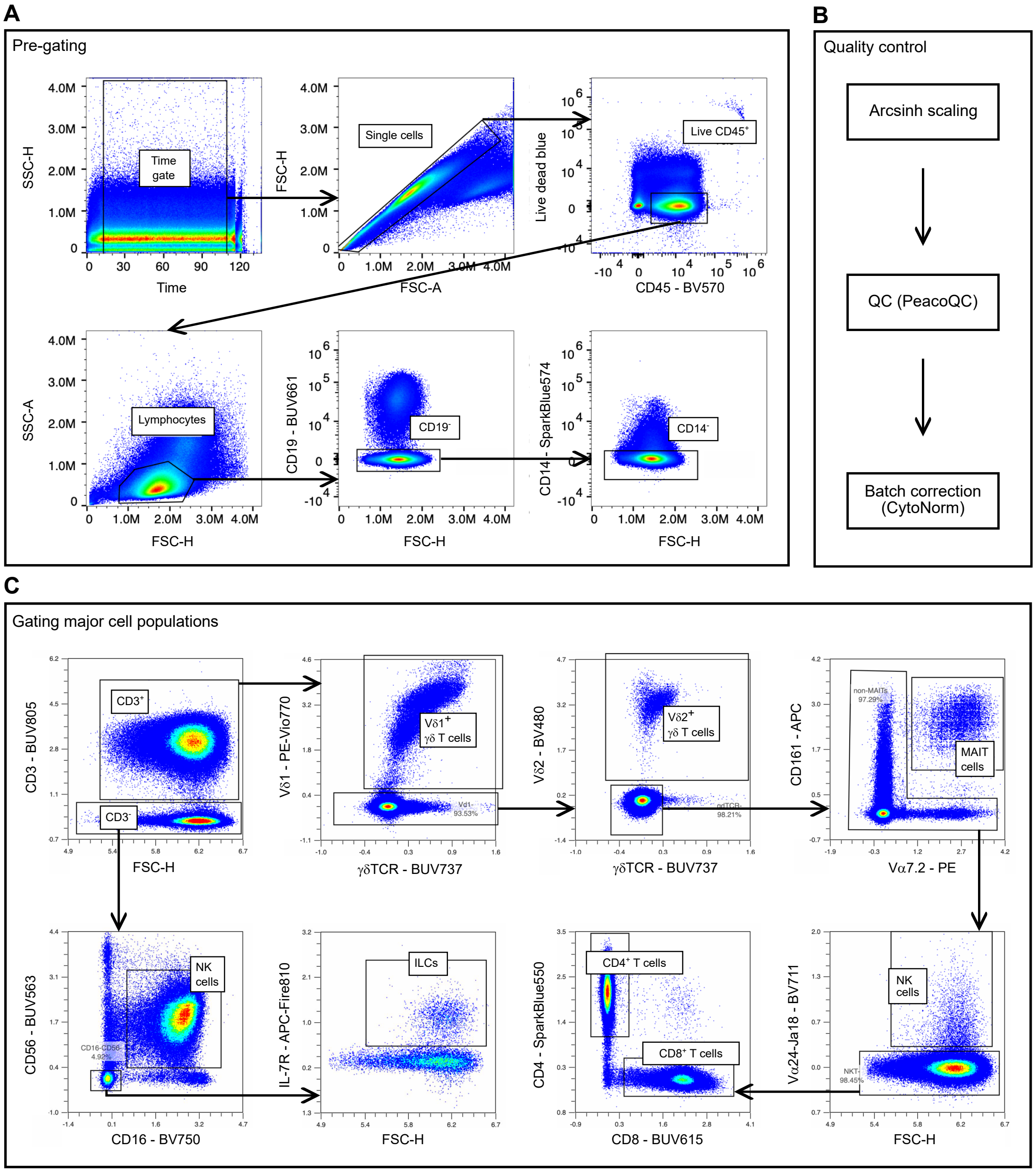


**Fig. S1. Immune cell phenotyping flow cytometry gating strategy.**
(**A**) Flow cytometry gating strategy for defining immune cell populations. (**B**) Samples were pre-gated, subjected to quality control (QC) including scaling and batch correction, then (**C**) gated for V$\delta$1$^+$ $\gamma\delta$ T cells, V$\delta$2$^+$ $\gamma\delta$ T cells, MAIT cells, invariant natural killer T (NKT) cells, conventional CD8$^+$ T cells (cytotoxic T lymphocytes, CTL), conventional CD4$^+$ T cells (T helper cells, Th), innate lymphoid cells (ILC) and natural killer (NK) cells.

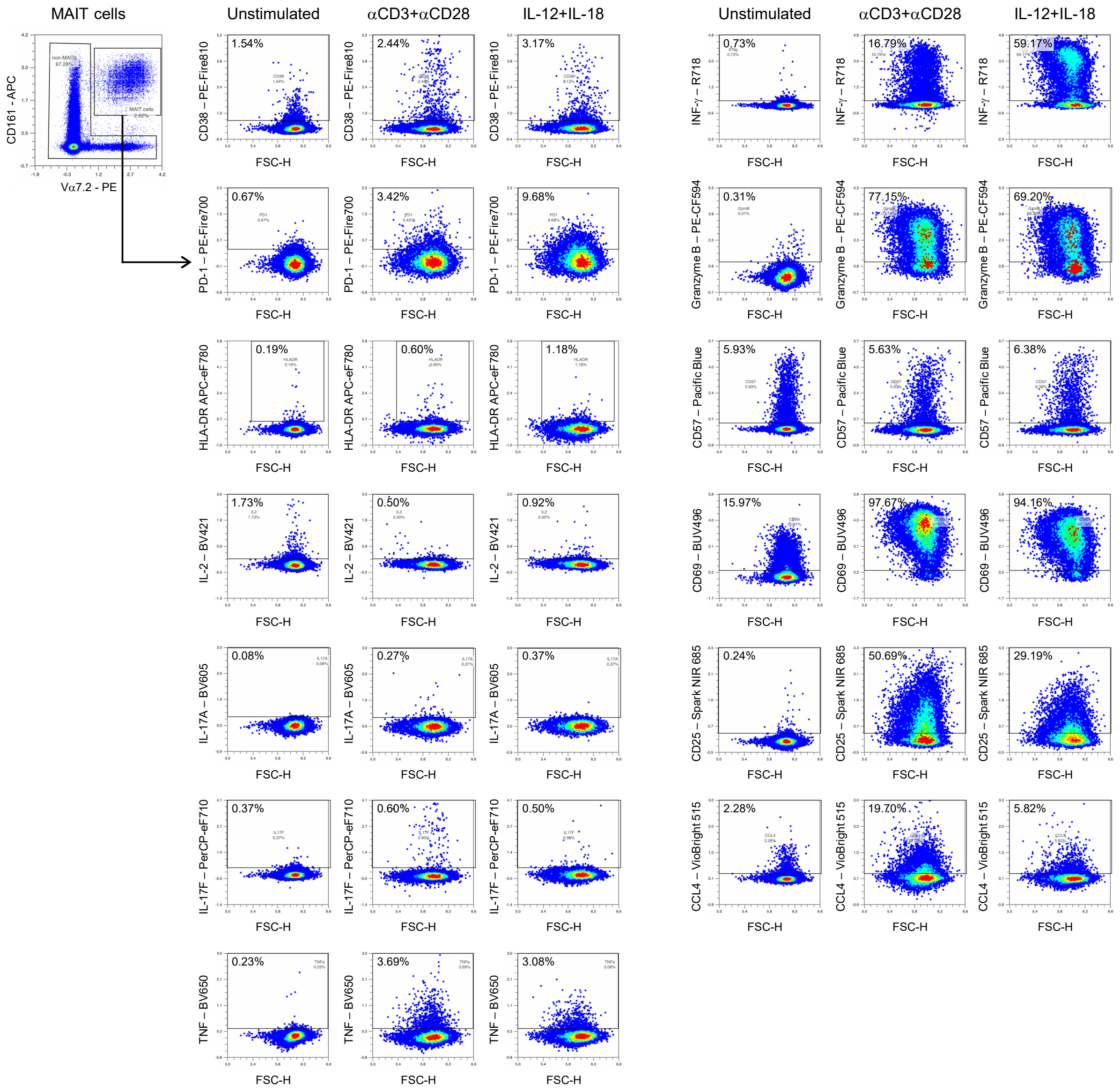


**Fig. S2. Activation marker flow cytometry gating strategy.**

Flow cytometry gating strategy for activation markers and cytokines on MAIT cells stimulated with media control ('unstimulated'), human monoclonal anti-CD3 and anti-CD28 antibodies or interleukin-12 (IL-12) and IL-18 for 24 h. Example plots from the same donor are shown.

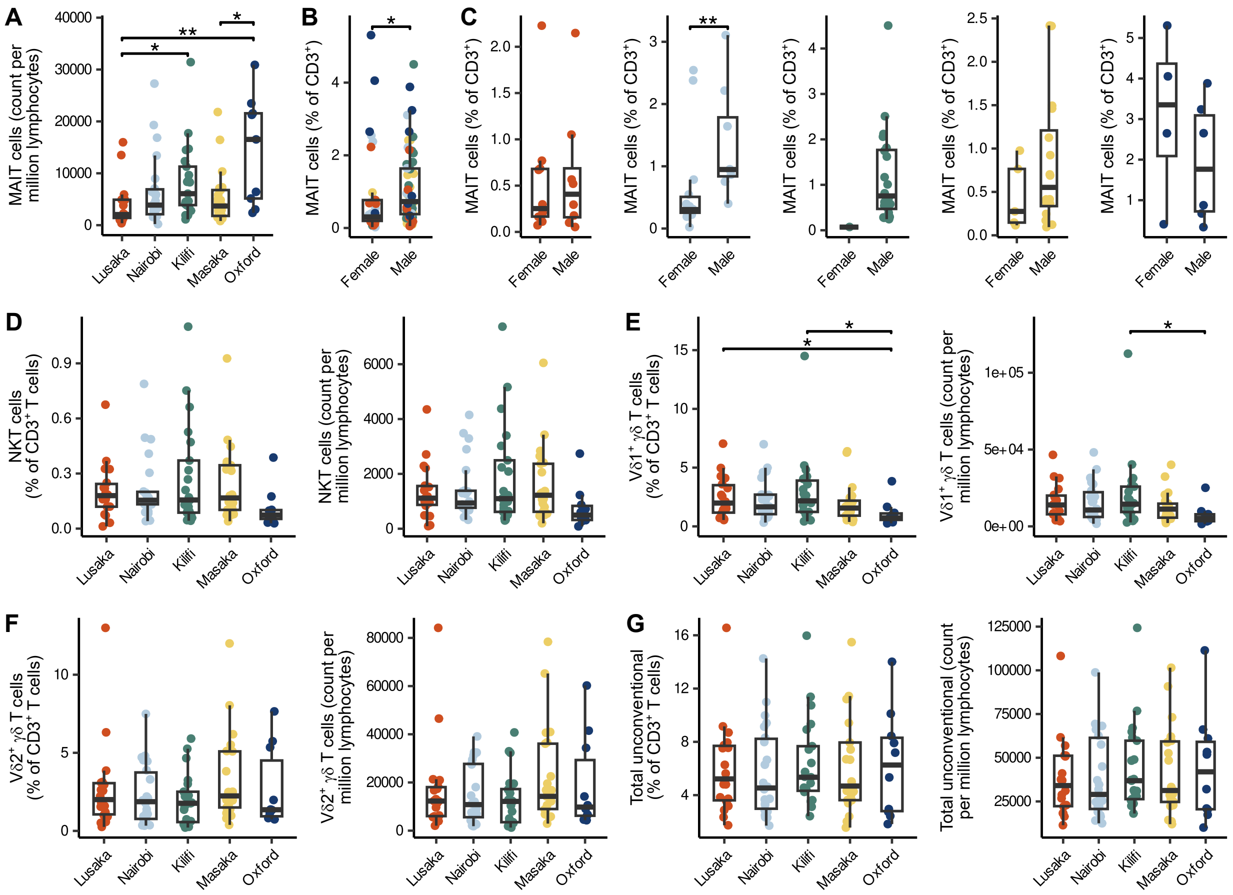


**Fig S3. Abundance and counts of unconventional T cells.**

(**A**) Absolute MAIT cell count (per million lymphocytes) by study site. (**B** and **C**), Abundance (percent of $CD3^+$ cells) of MAIT cells by sex across the (**B**) whole dataset and (**C**) in each individual study site. (**D** to **G**), Abundance (percent of $CD3^+$ cells) and absolute cell count (per million lymphocytes) of (**D**) NKT cells, (**E**) $V\delta1^+$ γδ T cells, (**F**) $V\delta2^+$ γδ T cells and (**G**) unconventional T cells (total of MAIT cells, $V\delta1^+$ γδ T cells, $V\delta2^+$ γδ T cells and NKT cells) by study site. All panels represent data from $n = 84$ biologically independent samples. Boxplots denote median, 25th and 75th percentiles, and whiskers to the largest value within 1.5 × interquartile range of the percentiles. Individual data points are shown. Significance determined by ANOVA with Tukey's post-hoc correction where assumptions met; otherwise, Kruskal-Wallis H test with Dunn's post-hoc test with Bonferroni's correction used. *, $p < 0.05$; **, $p < 0.01$; ***, $p < 0.001$ (adjusted p-values). For Figures [(**B**) and (**C**)], significance was determined by Mann Whitney U test.

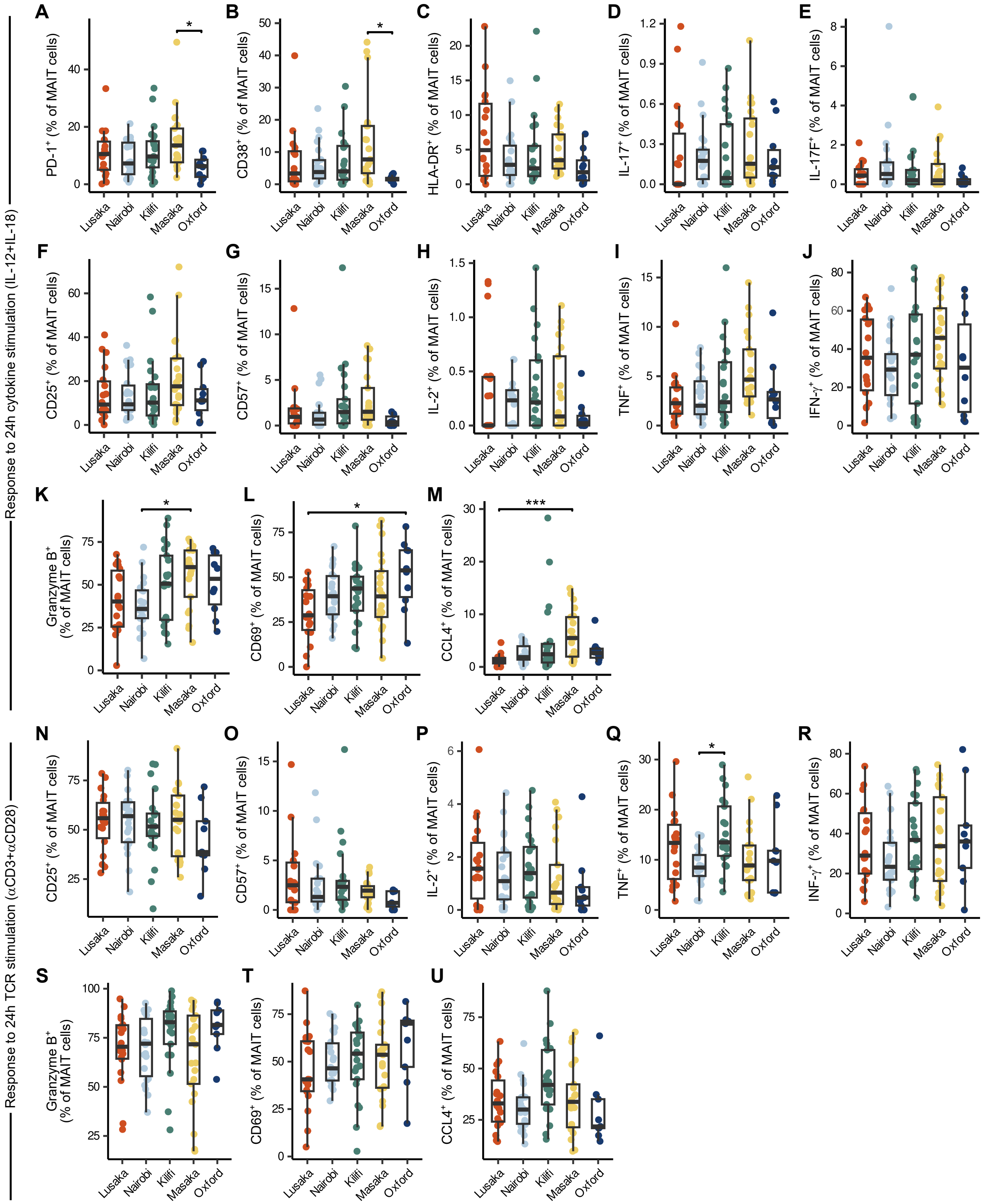


**Fig S4. Expression of markers following cytokine and TCR stimulation.**
(**A** to **M**) Abundance of (**A**) PD-1$^+$, (**B**) CD38$^+$, (**C**) HLA-DR$^+$, (**D**) IL-17A$^+$, (**E**) IL-17F$^+$, (**F**) CD25$^+$, (**G**), CD57$^+$, (**H**) IL-2$^+$, (**I**) TNF$^+$, (**J**) IFN-γ$^+$, (**K**) Granzyme B$^+$, (**L**) CD69$^+$, (**M**) CCL4$^+$ MAIT cells (change in percent expression) following 24-hour stimulation with interleukin (IL)-12

and IL-18 (cytokine stimulation) by study site. (**N** to **U**), Abundance of (**N**) $CD25^+$, (**O**), $CD57^+$, (**P**) IL-$2^+$, (**Q**) $TNF^+$, (**R**) IFN-$\gamma^+$, (**S**) Granzyme $B^+$, (**T**) $CD69^+$, (**U**) $CCL4^+$ MAIT cells (change in percent expression) following 24-hour stimulation with anti-CD3 and anti-CD28 monoclonal antibodies (TCR stimulation) by study site. All panels represent data from n = 84 biologically independent samples. Boxplots denote median, 25th and 75th percentiles, and whiskers to the largest value within 1.5 × interquartile range of the percentiles. Individual data points are shown. Significance determined by ANOVA with Tukey's post-hoc correction where assumptions met; otherwise, Kruskal-Wallis H test with Dunn's post-hoc test with Bonferroni's correction used. *, $p < 0.05$; **, $P < 0.01$; ***, $p < 0.001$ (adjusted p-values).

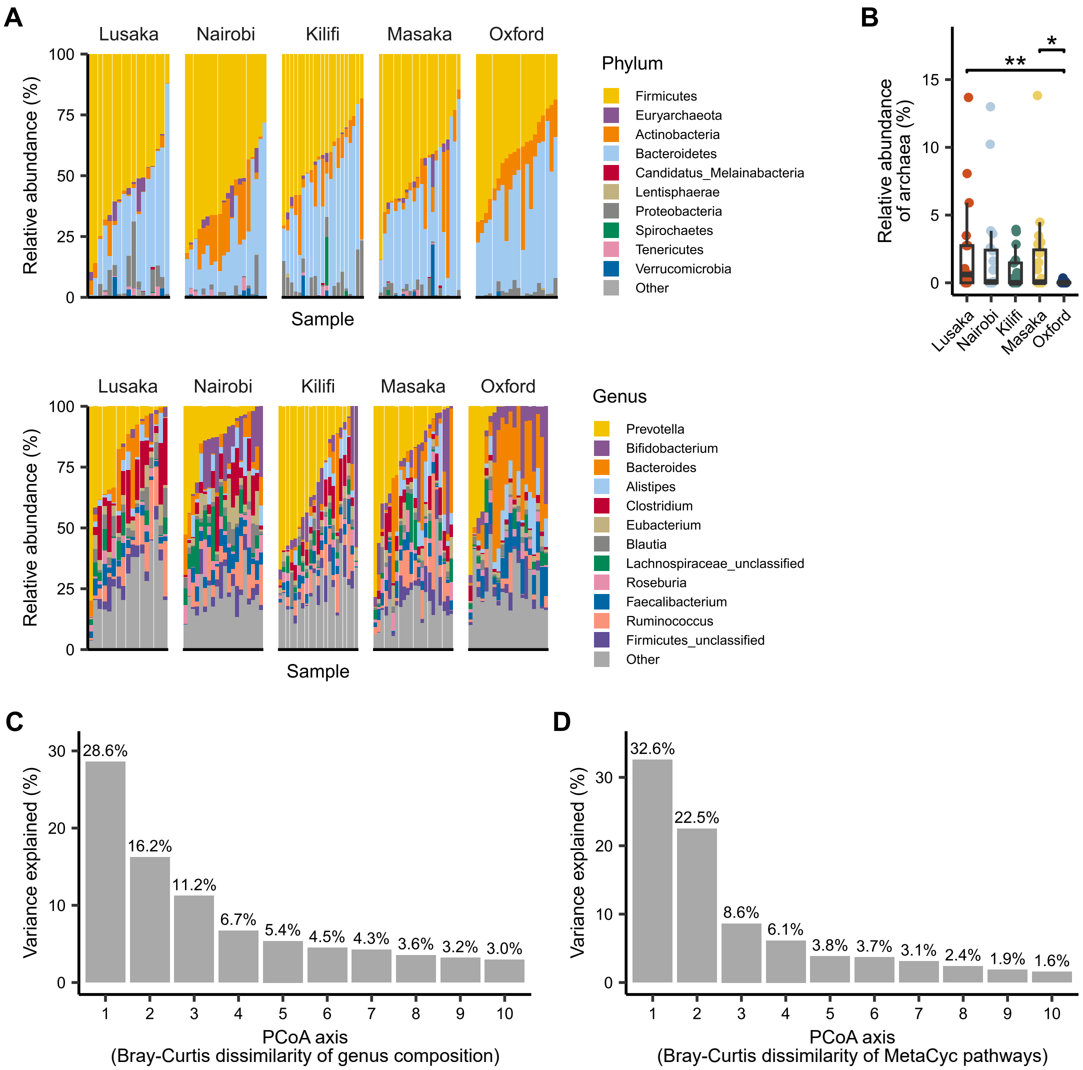


**Fig. S5. Microbiome composition and scree plots.**
(**A**) Relative abundance of the top 10 most abundant phyla and top 12 most abundant genera across individual microbiomes, separated by study site. Remaining taxa are grouped as "Other". (**B**) Relative abundance of archaea by study site. Significance determined by Kruskal-Wallis H test with Dunn's post-hoc test with Bonferroni's correction. *, $p < 0.05$; **, $p < 0.01$; ***, $p < 0.001$ (adjusted p-values). (**C** and **D**) Scree plot of variance explained by the first 10 principal coordinates of a Principal Coordinates Analysis (PCoA) performed on Bray-Curtis dissimilarity (BCD) on (**C**) genus-level relative abundances and (**D**) abundances of MetaCyc gene pathways.

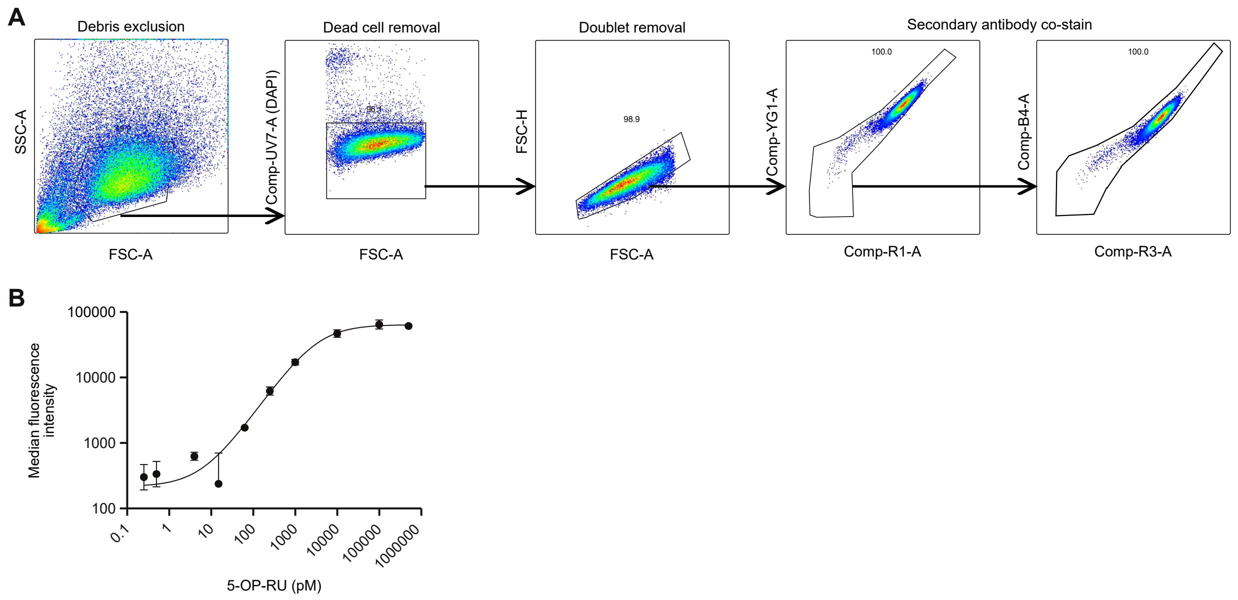


**Fig. S6. Flow cytometry gating strategy and dose-dependent 5-OP-RU response in THP-1 cells.**

(**A**) Gating strategy used for THP-1-MR1 cells. Debris were excluded based on FSC-A and SSC-A, dead cells were removed by DAPI staining, and cells were then gated for double positivity using the secondary antibodies, first on YG-1A versus R-1A and then on B4 versus R-3A. (**B**) Example of a 5-OP-RU standard curve for quantification in human faecal samples.

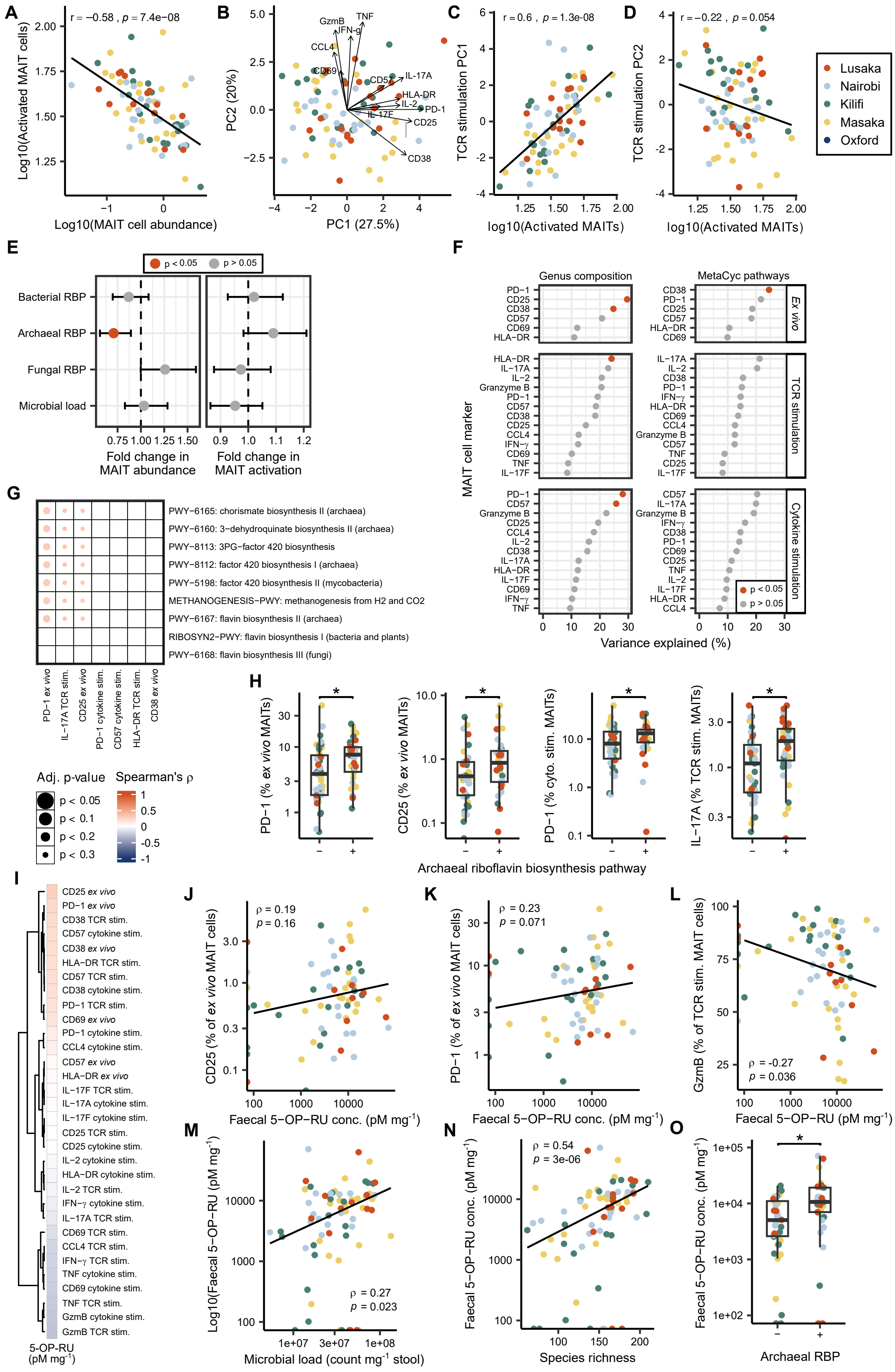

A
r = −0.58 , p = 7.4e−08
Log10(Activated MAIT cells)
Log10(MAIT cell abundance)
B
PC2 (20%)
PC1 (27.5%)
GzmB
IFN-g
TNF
CCL4
CD69
CD57
IL-17A
HLA-DR
IL-2
PD-1
IL-17F
CD25
CD38
C
r = 0.6 , p = 1.3e-08
TCR stimulation PC1
log10(Activated MAITs)
D
r = −0.22 , p = 0.054
TCR stimulation PC2
log10(Activated MAITs)
Lusaka
Nairobi
Kilifi
Masaka
Oxford
E
p < 0.05
p > 0.05
Bacterial RBP
Archaeal RBP
Fungal RBP
Microbial load
Fold change in MAIT abundance
Fold change in MAIT activation
F
Genus composition
MetaCyc pathways
MAIT cell marker
Ex vivo
TCR stimulation
Cytokine stimulation
Variance explained (%)
G
PWY−6165: chorismate biosynthesis II (archaea)
PWY−6160: 3−dehydroquinate biosynthesis II (archaea)
PWY−8113: 3PG−factor 420 biosynthesis
PWY−8112: factor 420 biosynthesis I (archaea)
PWY−5198: factor 420 biosynthesis II (mycobacteria)
METHANOGENESIS−PWY: methanogenesis from H2 and CO2
PWY−6167: flavin biosynthesis II (archaea)
RIBOSYN2−PWY: flavin biosynthesis I (bacteria and plants)
PWY−6168: flavin biosynthesis III (fungi)
PD−1 ex vivo
IL−17A TCR stim.
CD25 ex vivo
PD−1 cytokine stim.
CD57 cytokine stim.
HLA−DR TCR stim.
CD38 ex vivo
Adj. p-value
p < 0.05
p < 0.1
p < 0.2
p < 0.3
Spearman's ρ
H
PD−1 (% ex vivo MAITs)
CD25 (% ex vivo MAITs)
PD−1 (% cyto. stim. MAITs)
IL−17A (% TCR stim. MAITs)
Archaeal riboflavin biosynthesis pathway
I
CD25 ex vivo
PD−1 ex vivo
CD38 TCR stim.
CD57 cytokine stim.
CD38 ex vivo
HLA−DR TCR stim.
CD57 TCR stim.
CD38 cytokine stim.
PD−1 TCR stim.
CD69 ex vivo
PD−1 cytokine stim.
CCL4 cytokine stim.
CD57 ex vivo
HLA−DR ex vivo
IL−17F TCR stim.
IL−17A cytokine stim.
IL−17F cytokine stim.
CD25 TCR stim.
CD25 cytokine stim.
IL−2 cytokine stim.
HLA−DR cytokine stim.
IL−2 TCR stim.
IFN−γ cytokine stim.
IL−17A TCR stim.
CD69 TCR stim.
CCL4 TCR stim.
IFN−γ TCR stim.
TNF cytokine stim.
CD69 cytokine stim.
TNF TCR stim.
GzmB cytokine stim.
GzmB TCR stim.
5-OP-RU (pM mg-1)
J
ρ = 0.19
p = 0.16
CD25 (% of ex vivo MAIT cells)
Faecal 5−OP−RU conc. (pM mg-1)
K
ρ = 0.23
p = 0.071
PD−1 (% of ex vivo MAIT cells)
Faecal 5−OP−RU conc. (pM mg-1)
L
ρ = -0.27
p = 0.036
GzmB (% of TCR stim. MAIT cells)
Faecal 5−OP−RU (pM mg-1)
M
ρ = 0.27
p = 0.023
Log10(Faecal 5−OP−RU (pM mg-1)
Microbial load (count mg-1 stool)
N
ρ = 0.54
p = 3e−06
Faecal 5−OP−RU conc. (pM mg-1)
Species richness
O
Faecal 5−OP−RU conc. (pM mg-1)
Archaeal RBP

**Fig. S7. Key analyses repeated with UK population excluded.**
(**A**) Log10-transformed MAIT abundance (percent of $CD3^+$ T cells) by log10-transformed abundance of activated MAIT cells. (**B**) Principal component analysis (PCA) on z-score normalised change in percent expression of markers by MAIT cells following TCR stimulation. Each data point represents an individual participant, coloured by study site. Arrows represent loadings of each marker. (**C** and **D**) Log10-transformed abundance of activated MAIT cells (percent of MAIT cells) by (**C**) PC1 and (**D**) PC2. (**E**) Coefficient plot of microbiome predictors of log-transformed MAIT abundance and log-transformed abundance of activated MAIT cells. Points indicate estimated regression coefficients (β) from a linear model including z-scaled abundance of the bacterial, fungal and archaeal riboflavin synthesis pathways, z-scaled microbial load, sex and age with study site as a random effect. Bars indicate 95% confidence intervals. Colour denotes significance ($p < 0.05$) by type III F-tests with Satterthwaite's approximation. (**F**) Variance explained (R-squared) from linear models (MAIT cell phenotypic marker ~ PC1 + … + PC10), where PC1-PC10 are principal coordinates from a PCoA on genus- or pathway-level BCD. Colour denotes significance ($P < 0.05$) by overall model F-test. (**G**) Spearman's rank correlations between MetaCyc gene pathways and MAIT cell phenotypic markers. Colour indicates Spearman's ρ; circle size denotes Benjamini-Hochberg-adjusted p-values. (**H**) Abundance of MAIT cells expressing phenotypic markers (ex vivo or following cytokine or TCR stimulation) in individuals where the archaeal riboflavin biosynthesis pathway (RBP) is detectable (+) or undetectable (-) by shotgun metagenomic sequencing. Significance determined by Mann-Whitney U test. (**I**) Heatmap of Spearman's rank correlations between faecal 5-OP-RU concentrations and MAIT cell phenotypic or functional parameters measured directly ex vivo or following stimulation. (**J** to **L**) Correlation between faecal 5-OP-RU concentration and circulating MAIT cell expression of (J) CD25 and (K) PD-1 directly ex vivo and (L) granzyme B following TCR stimulation. (**M** and **N**) Correlation between faecal 5-OP-RU concentration and (M) faecal microbial load and (**N**) faecal microbiome species richness. (**O**) Faecal 5-OP-RU concentration in individuals with the archaeal riboflavin biosynthesis pathway (RBP) detectable (+) or undetectable (−) by shotgun metagenomic sequencing. For scatter plots, r denotes Pearson's correlation and ρ denotes Spearman's correlation coefficient. Data points are coloured by study site. Line represents the linear regression fit (least-squares line) across all data points. For boxplots comparing two groups, significance determined by Mann-Whitney U test.

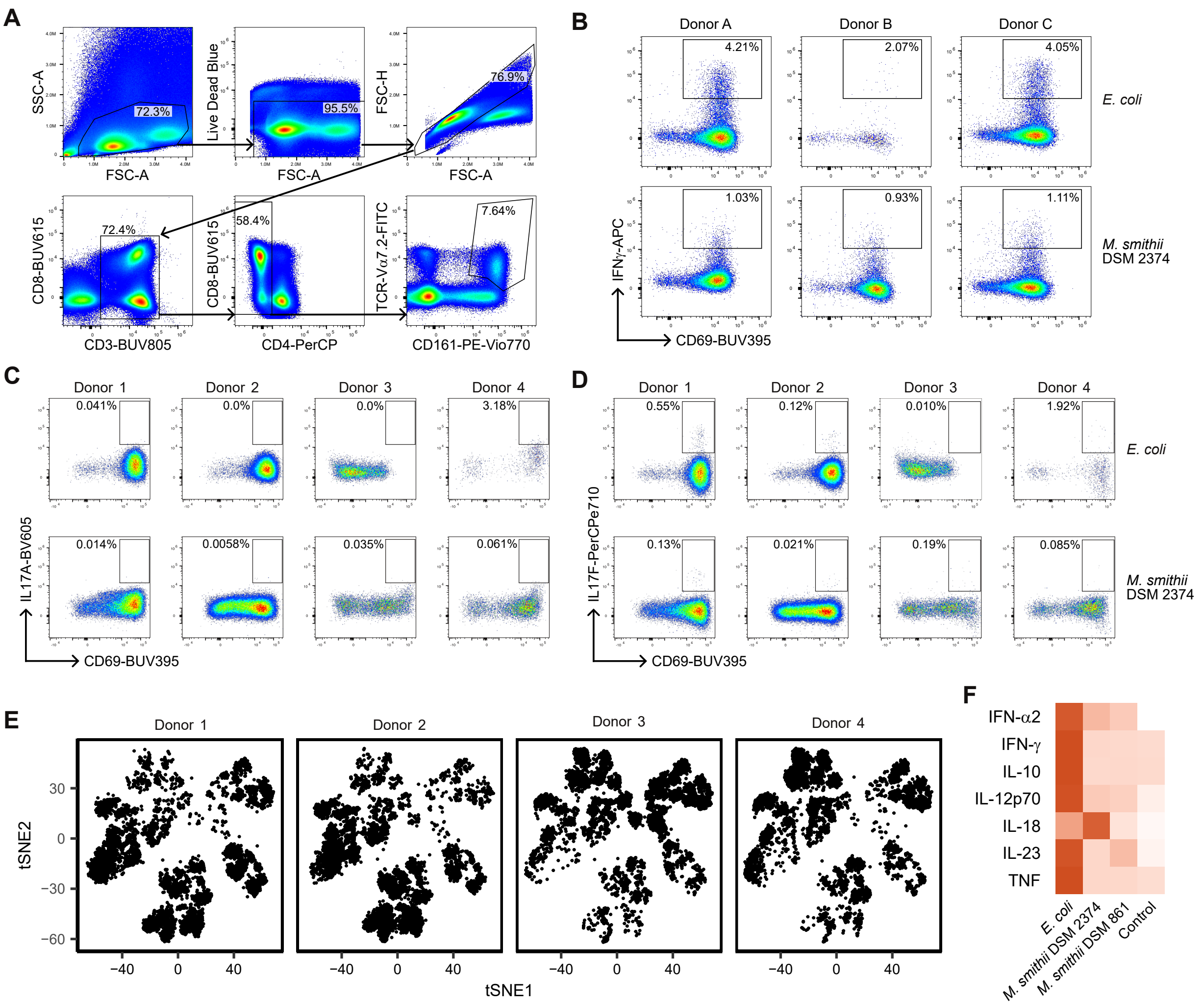


**Fig. S8. Flow cytometry gating strategy and supporting data for Figure 4.**
(**A**) Gating strategy for MAIT cells: lymphocyte-sized cells were first selected based on forward- and side-scatter characteristics, followed by exclusion of dead cells using the LD Blue viability dye and selection of singlets. $CD3^+$ T cells were then identified, and $CD4^-$ cells were selected. MAIT cells were defined as $CD161^{hi}$ TCR-Vα$7.2^+$ cells. (**B** to **D**) CD69 and IFN-γ (B) IL-17A (**C**) and IL-17F (D) expression in MAIT cells after activation with E. coli or M. smithii. (**E**) Donor distribution in t-SNE space. (**F**) Heatmap showing cytokine concentrations in supernatants after microbial stimulation, corresponding to Fig. 4H, in the presence of an isotype control antibody.

### Table S1. Human study population demographics.

| Trial | Study site | n | Female n (%) | Age Median (range) | Demographic |
|---|---|---|---|---|---|
| HIV-CORE 005.1 | Oxford, UK | 10 | 2 (20%) | 24.5 (21-63) | Healthy adults recruited in Oxford at low likelihood of acquiring HIV-1. |
| HIV-CORE 005.2 | Oxford, UK | 10 | 4 (40%) | 30 (26-63) | Healthy adults recruited in Oxford at low likelihood of acquiring HIV-1. |
| HIV-CORE 006 | Lusaka, Zambia | 22 | 11 (50%) | 34 (24-47) | Mostly within 5-10 km of study site (3 km north of Lusaka), though some participants from peri-urban communities outside this radius. |
| HIV-CORE 006 | Nairobi, Kenya | 22 | 14 (63.5%) | 27 (19-37) | Population of low-income earners living in informal settlements, who frequently migrate. |
| HIV-CORE 006 | Kilifi, Kenya | 22 | 1 (4.55%) | 32 (27-50) | Gay males and females, people who inject drugs, low-income women, sex workers and young men who have sex with men. All participants on PreP. |
| HIV-CORE 006 | Masaka, Uganda | 22 | 5 (22.7%) | 29.5 (19-39) | Within 30 km radius of Masaka; mainly agricultural population. |

**Table S2. Antibody reagent table.**
Experiment 1 refers to human population phenotyping in Figure 1; Experiment 2 refers to in vitro MAIT cell activation assays in Figure 4.

| Excitation laser | Fluorochrome | Marker | Clone | Manufacturer | Cat. no. | RRID | Dilution (1 in X) | Experiment |
|---|---|---|---|---|---|---|---|---|
| 355nm/UV | BUV395 | CD45RA | HI100 | BD Biosciences | 740298 | AB_2740037 | 200 | 1 |
| 355nm/UV | BUV496 | CD69 | FN50 | BD Biosciences | 750214 | AB_2874415 | 100 | 1 |
| 355nm/UV | BUV563 | CD56 | NCAM16.2 | BD Biosciences | 612928 | AB_2870213 | 200 | 1 |
| 355nm/UV | BUV615 | CD8 | SK1 | BD Biosciences | 612995 | AB_2870266 | 200 | 1 & 2 |
| 355nm/UV | BUV661 | CD19 | HIB19 | BD Biosciences | 741604 | AB_2871012 | 200 | 1 |
| 355nm/UV | BUV737 | γδ TCR | 11F2 | BD Biosciences | 748533 | AB_2872944 | 200 | 1 |
| 355nm/UV | BUV805 | CD3 | UCHT1 | BD Biosciences | 612895 | AB_2870183 | 25 | 1 & 2 |
| 405nm/V | BV421 | IL2 | MQ1-17H12 | BioLegend | 500327 | AB_10897949 | 200 | 1 |
| 405nm/V | Pacific Blue™ | CD57 | HNK-1 | BioLegend | 359607 | AB_2562458 | 200 | 1 |
| 405nm/V | BV480 | Vδ2 TCR | B6 | BD Biosciences | 746567 | AB_2743853 | 100 | 1 |
| 405nm/V | Brilliant Violet 570™ | CD45 | HI30 | BioLegend | 304033 | AB_10899568 | 200 | 1 |
| 405nm/V | Brilliant Violet 605™ | IL-17A | BL168 | BioLegend | 512325 | AB_11218595 | 200 | 1 & 2 |
| 405nm/V | Brilliant Violet 650™ | TNF | MAb11 | BioLegend | 502937 | AB_2561355 | 200 | 1 & 2 |
| 405nm/V | Brilliant Violet 711™ | TCR Vα24-Jα18 | 6B11 | BioLegend | 342921 | AB_2572067 | 200 | 1 |
| 405nm/V | BV750 | CD16 | 3G8 | BD Biosciences | 747461 | AB_2872137 | 200 | 1 |
| 405nm/V | Brilliant Violet 785™ | CD197 (CCR7) | G043H7 | BioLegend | 353229 | AB_2561371 | 25 | 1 |
| 488nm/B | Vio® B515 | CCL4 (MIP-1β) | REA511 | Miltenyi Biotec | 130-129-199 | AB_2922004 | 100 | 1 |
| 488nm/B | Spark Blue™ 550 | CD4 | SK3 | BioLegend | 344655 | AB_2819978 | 200 | 1 & 2 |
| 488nm/B | Spark Blue™ 574 | CD14 | HCD14 | BioLegend | 325635 | AB_2904346 | 200 | 1 |
| 488nm/B | PerCP-eFluor™ 710 | IL-17F | SHLR17 | Thermo Fisher Scientific | 46-7169-42 | AB_10596829 | 100 | 1 & 2 |
| 561nm/YG | PE | TCR Vα7.2 | 3C10 | BioLegend | 351705 | AB_10933262 | 100 | 1 |
| 561nm/YG | PE-CF594 | Granzyme B | GB11 | BD Biosciences | 562462 | AB_2737618 | 200 | 1 & 2 |
| 561nm/YG | PE/Fire™ 700 | CD279 (PD-1) | A17188B | BioLegend | 621622 | AB_2910489 | 100 | 1 & 2 |
| 561nm/YG | PE-Vio® 770 | TCR Vδ1 | REA173 | Miltenyi Biotec | 130-117-809 | AB_2751427 | 100 | 1 |
| 561nm/YG | PE/Fire™ 810 | CD38 | S17015F | BioLegend | 397225 | AB_2894562 | 200 | 1 & 2 |
| 640nm/R | APC | CD161 | 191B8 | Miltenyi Biotec | 130-114-116 | AB_2733345 | 200 | 1 |
| 640nm/R | Spark NIR™ 685 | CD25 | M-A251 | BioLegend | 356151 | AB_2888804 | 200 | 1 |
| 640nm/R | R718 | IFN-γ | B27 | BD Biosciences | 566960 | AB_2869972 | 200 | 1 & 2 |
| 640nm/R | APC-eFluor™ 780 | HLA-DR | L243 | Invitrogen | 47-9952-42 | AB_2784721 | 200 | 1 |
| 640nm/R | APC/Fire™ 810 | CD127 (IL-7Rα) | A019D5 | BioLegend | 351373 | AB_2904371 | 200 | 1 |
| 355nm/UV | BUV395 | CD69 | FN50 | BD Biosciences | 564364 | AB_3106238 | 200 | 2 |
| 405nm/V | BV421 | CD25 | BC96 | BioLegend | 302630 | AB_10896914 | 70 | 2 |
| 405nm/V | BV785 | HLA-DR | L243 | BioLegend | 307642 | AB_2561360 | 150 | 2 |
| 488nm/B | FITC | TCR Vα7.2 | REA179 | Miltenyi Biotec | 130-123-685 | AB_2811544 | 150 | 2 |
| 488nm/B | PerCP | CD4 | SK3 | BioLegend | 344624 | AB_2563325 | 100 | 2 |
| 561nm/YG | PE-Vio® 770 | CD161 | REA631 | Miltenyi Biotec | 130-113-597 | AB_2726190 | 150 | 2 |
| 640nm/R | APC | CD137 | REA765 | Miltenyi Biotec | 130-110-764 | AB_2654988 | 100 | 2 |
